# Targeting SCD in SCD-amplified prostate cancer inhibits growth in bone by modulating cellular stress, mTOR, and DNA damage pathways

**DOI:** 10.1101/2025.06.09.658523

**Authors:** Alexis Wilson, Mackenzie Herroon, Shane Mecca, Laimar Garmo, Jacob Lindquist, Shrila Rajendran, Steve M. Patrick, Izabela Podgorski

**Affiliations:** Department of Pharmacology, Wayne State University School of Medicine, Detroit, MI, USA; Karmanos Cancer Institute, Wayne State University School of Medicine, Detroit, MI, USA

**Author notes:** Corresponding Author: Izabela Podgorski Wayne State University School of Medicine Department of Pharmacology 540 E. Canfield, Rm 6312 Detroit, MI 48201. The authors disclose no potential conflicts of interest.

**Keywords:** Bone marrow adipocytes, bone metastasis, prostate cancer, lipid desaturation, ER stress, DNA damage

## Abstract

The bone microenvironment is abundant in adipocytes and fosters metastatic progression, but the underlying mechanisms are not fully understood. We hypothesize that Stearoyl-Coenzyme A Desaturase (SCD) acts as a tumor-promoting enzyme by modulating cellular stress to support the growth and survival of prostate cancer (PCa) in bone. We observe that SCD-amplified PCa cells are highly sensitive to SCD loss and show reduced PCa spheroid size, diminished mTOR signaling, and increased ER stress. Notably, SCD expression is further increased by adipocytes in SCD-amplified cell lines, and its loss increases DNA damage and activates repair pathways in PCa cells only when exposed to adipocytes. Furthermore, we observe PCa cell lines utilize SCD to regulate adipocyte-induced lipid peroxidation. Aligned with these results, pharmacological SCD inhibition in mice bearing SCD-amplified bone tumors reduces tumor size and reveals histochemical evidence of increased ER stress and DNA damage. Collectively, our data highlight the impact of SCD loss on SCD-amplified tumors and suggest germline characteristics of tumors may dictate their response to redox insult and the possibility of targeting DNA repair pathways in combination with SCD inhibition.

## 1. INTRODUCTION

Alterations in lipid uptake and metabolism have been recognized as some of the most prominent adaptive mechanisms in cancer, contributing to proliferation and rapid tumor growth [1]. Certain lipid species are particularly advantageous for cancer cells and play tumor-supportive roles in promoting growth, survival, and progression. Monounsaturated fatty acids (MUFAs), specifically, are known to make up cell membrane composition and play role(s) in reducing membrane sensitivity to polyunsaturated fatty acid (PUFA)-dependent oxidation [2]. Higher MUFA levels have been found in cancerous tissues and have been shown to modulate tumorigenic pathways, such as activating PI3K and ERK signaling via G-coupled protein receptor 40 [3]. Stearoyl coenzyme A desaturase (SCD) is the primary endoplasmic reticulum (ER)-resident enzyme involved in the conversion of saturated fatty acids (SFAs) to MUFAs. We have reported SCD gene amplification in metastatic prostate cancer, and others have demonstrated increased SCD expression across several other cancer types [4, 5]. SCD has been suggested to play critical roles in tumor formation, progression, and survival of cancer cells [6]. This underscores SCD activity as a possible major contributor to tumor progression, especially in harsh microenvironments abundant in lipids, such as bone.

Bone is the primary site for metastatic prostate cancer (PCa), and its ability to adapt and thrive in the lipid-rich bone marrow poses a major clinical challenge. Adipocytes constitute a major cell type in bone, and previous studies from our laboratory and others have shown that tumor cells disseminated to or originating from bone promote lipolytic behavior in marrow adipocytes, which then release lipids that are absorbed by the tumor cells [4, 7-10]. We previously reported that adipocyte-supplied lipids disrupt redox balance and induce cellular stress pathways in PCa cells, yet allow them to survive and progress within the harsh bone metastatic niche [11, 12]. SCD expression has been shown to be induced by SFAs, possibly as a mechanism to support ER homeostasis, prevent membrane oxidation, and promote a cellular adaptive response to stress-inducing lipid species [5]. However, the molecular mechanisms that drive this adaptive response are poorly understood.

The present study aimed to investigate the molecular mechanisms by which SCD promotes the adaptive response to stress in metastatic PCa. Using *in vitro* models, we investigated the protein expression of SCD in various PCa cell lines and utilized 3D culture methods to determine if the expression levels of SCD correlated with PCa growth and survival during SCD loss. RNAseq analyses and functional assays were used to examine the molecular pathways dysregulated during SCD deficiency. *In vivo* methods included intratibial experimental bone tumors to evaluate the effect of SCD inhibition on PCa growth and survival. Collectively, our studies explored a novel molecular mechanism involving SCD in tumor cell adaptive response to stress in the bone metastatic niche, which has therapeutic implications for bone metastatic PCa and other cancers that reside in skeletal sites.

## 2. MATERIALS AND METHODS

### 2.1 Materials

DMEM, RPMI-1640, MEMα, insulin, and other chemicals, unless otherwise stated, were obtained from Sigma-Aldrich (St. Louis, MO). HyClone FBS, TaqMan reagents, and RNAiMAX were from ThermoFisher Scientific (Waltham, MA). Trypsin-EDTA was from Invitrogen (Carlsbad, CA). PureCol® collagen type I was from Advanced Biomatrix (San Diego, CA). Transwell cell-support systems were from Corning (Corning, NY). β-tubulin (#E7-C) antibody was from Developmental Studies Hybridoma Bank (Iowa City, IA). Antibodies to HSPA5 (BIP; #3177), ATF4 (#11815), γ-H2AX (#9718), p-AKT (#4060), Total AKT (#9272), p-p70S6K (#9205), Total p70S6K (#34475), p-PRAS40 (#13175), Total PRAS40 (#2691), p-NDRG1 (#5482), Total NDRG1 (#9485), and p-4EBP1 (#9459) were from Cell Signaling Technology (Danvers, MA). Antibody to SCD (#28678-1-AP) was from Proteintech (Rosemont, IL) and Bioss (#3787R). StemXVivo Adipogenic Supplement and Cultrex were from R&D Systems (Minneapolis, MN). RNeasy Mini Kits were from Qiagen (Valencia, CA). QuantSeq 3’ mRNA-Seq Library Prep Kit FWD were from Lexogen (Vienna, Austria). Luminata Forte Western HRP substrate was from MilliporeSigma (Burlington, MA). Rosiglitazone was from Cayman Chemical (Ann Arbor, MI).

### 2.2 Cell Lines

PC3, 22Rv1, and C42B cells were purchased from ATCC (Manassas, VA). ARCaP(M) cells were purchased from Novicure Biotechnology (Birmingham, AL). PC3 and C42B cells were cultured in DMEM with 10% FBS, ARCaP(M) cells were cultured in RPMI-1640 medium with 5% FBS, and 22Rv1 cells were cultured with RPM1-1640 with 10% FBS. All media were supplemented with 25mM HEPES, and 100U/ml penicillin-streptomycin. Primary mouse bone marrow stromal cells (mBMSC) were isolated from tibiae and femurs of 6- to 8-week old FVB/N mice and differentiated into adipocytes according to previously established protocols [8, 13]. The human cell lines used in this study were authenticated by the WSU Genomics facility. All cell lines are routinely tested for mycoplasma using LookOut Mycoplasma PCR Detection Kit (Sigma).

### 2.3 Animals

All experiments involving mice were performed in accordance with the protocol approved by the Institutional Animal Investigational Committee of Wayne State University (IACUC-21-12-4269). All animal experiments complied with the ARRIVE guidelines and followed the National Research Council’s Guide for the Care of Use of Laboratory Animals. *In vivo*, xenograft studies were performed in 8- to 10-week-old male mice in the FVB/N background with a homozygous null mutation in the Rag-1 gene (FVB/N, Rag-1^−/−^). All mice were bred in-house.

### 2.4 Pharmacological SCD and mTOR inhibition in Transwell co-culture

The mBMSC cells were embedded in Collagen I, plated in 6-well plates, and differentiated into adipocytes according to our previously published protocols [14, 8]. PC3, ARCaP(M), C42B, or 22Rv1 cells were seeded in 6-well plates or Transwell filters and cultured overnight. SCD inhibitor treatment with 1µM CAY10566 (Cayman Chemical, Ann Arbor, MI; concentrations determined experimentally) or 250µM A-939572 (Cayman Chemical, Ann Arbor, MI; concentrations determined experimentally), or mTORC1 inhibitor treatment with Everolimus (EVO) (Cayman Chemical, Ann Arbor, MI; concentrations determined experimentally) at 10nM or 50nM was applied the next day upon merging adipocyte and tumor cell cultures into the Transwell system. After 48 hours of co-culture, tumor cells were collected and processed for RNA extraction and protein analysis as we described [14, 8] [14, 7, 11, 12].

### 2.5 siRNA Approaches

Tumor cells were seeded on 6-well plates or on Transwell filters and grown overnight. The following day, a unique 27mer siRNA duplex targeting SCD transcripts (OriGene-SR321692) or Trilencer-27 Universal scrambled negative control (Origene-SR30002) was added using RNAiMAX transfection reagent at a final concentration of 25pmol (based on the manufacturer’s protocol). After 6 hours, the media was replaced, and Transwell filters containing transfected tumor cells were merged with bone marrow adipocytes. After 48 hours, cells were collected and processed for RNA and protein analyses as described above. Three unique 27mer siRNA duplexes that efficiently knocked down SCD transcripts were used.

### 2.6 TaqMan RT PCR analyses

The cDNA was prepared from 1-2μg of total RNA using High-Capacity cDNA Reverse Transcription kit (Applied Biosystems). Gene expression analyses were performed using TaqMan^®^ Individual Gene Expression assays for Human *SCD* (Hs01682761), *FERMT1* (Hs00916793), *GADD45A* (Hs00169255), *SOD2* (Hs00167309), *CD36* (Hs01567185), and *FABP4* (Hs010086177). Assays were conducted on at least three biological replicates using TaqMan^®^ Fast Universal PCR Master Mix and 50ng of cDNA/well, and all reactions were run on an Applied Biosystems StepOnePlus™ system. All genes were normalized to *hypoxanthine phosphoribosyltransferase* (*HPRT1*; Hs02800695). DataAssist™ Software (Applied Biosystems) was used for all analyses.

### 2.7 Immunoblotting

Lysate and media samples were loaded based on DNA/protein concentrations and the corresponding lysates were electrophoresed on Novex WedgeWell 4-20% Tris-Glycine Gels (Invitrogen), transferred to PVDF membranes and immunoblotted for SCD, HSPA5 (BIP), ATF4, p-AKT, Total AKT, p-p70S6K, Total p70S6K, p-PRAS40, Total PRAS40, p-NDRG1, Total NDRG1, and p-4EBP1, all at 1:1000. All horseradish peroxidase-labeled secondary antibodies were used at 1:10,000. Quantification and analyses of bands were performed using Fiji 2 (ImageJ, NIH, Bethesda, Maryland).

### 2.8 Lipid peroxidation assay

PC3 or ARCaP(M) cells were seeded on glass coverslips in a 24-well plate at a density of 50,000/coverslip overnight and then transferred to the top of a Transwell filter over adipocytes (Transwell) or into an empty 6-well plate (Control) (2 coverslips/well) and cultured for 24 or 48 hours. For SCD inhibition experiments, cells were treated with 1µM CAY10566 or siRNA against SCD as described above. On the day of imaging, cells were incubated with 10μM BODIPY 581/591 C11 (ThermoFisher, D3861) for 30 minutes at 37°C in 5% CO_2_. Following three washes with Live Cell Imaging Solution, pH 7.4 (Life Technologies Corporation, Eugene, DR), images (excitation of 581/488 nm and emission of 591/510 nm) were captured with a Zeiss LSM 780 confocal microscope (Carl Zeiss AG, Gottingen, Germany) using a 40× immersion lens. Integrated density of green (510nm) fluorescence/cell number, indicative of probe oxidation, was calculated using Fiji 2.

### 2.9 Immunofluorescence analyses

For assessment of lipid droplets in response to adipocyte exposure, PC3 or ARCaP(M) cells were grown in Transwell co-culture for 48 hrs as described above and stained with BODIPY 493/503 (1:1000) (Thermofisher, D3922) for 1 hour after fixing with 3.7% formaldehyde. Nuclei were stained by Hoechst dye (Invitrogen). Following three washes with Live Cell Imaging Solution, images were captured with a Zeiss LSM 780 confocal microscope using a 40× immersion lens. Lipid droplet count/cell number was calculated using the analyze particle function using Fiji 2. For γ-H2AX, ARCaP(M), PC3, 22Rv1, or C42B cells were grown on glass coverslips under control or Transwell conditions in the presence or absence of 1μM CAY10566 for 48 hours. Cells were fixed with 3.7% formaldehyde and stained with anti-γ-H2AX antibody (1:100) at 4°C overnight. Alexa Fluor 488-conjugated goat anti-rabbit IgG (1:1000) was used as a secondary antibody, and DAPI was used as a nuclear stain. Coverslips were mounted with Vectashield (Vector Laboratories, Newark, CA) and imaged on a Zeiss LSM 780 confocal microscope using a 63× oil immersion lens.

### 2.10 Alkaline Comet Assay

An alkaline comet assay was used to analyze DNA damage upon SCD inhibition using CAY10566. ARCaP(M) cells in alone conditions or Transwell co-culture were treated with 0.1% DMSO or 1μM CAY10566 as described above. ∼20,000 cells were embedded in 0.5% low-melting agarose (Fisher Scientific, BP165-25) and spread on slides coated with 1.5% Standard Low – mr Agarose (BioRad, 162-0100) and allowed to solidify. Slides were placed into 4°C lysis buffer (2.5M NaCl, 100mM EDTA, 10mM Tris Base, 1% Triton X-100, pH 10) for 1 hour. Slides were removed from the lysis buffer and placed in the electrophoresis tank with 4°C electrophoresis buffer (0.3M NaOH, 1mM EDTA) to incubate for 20 minutes, and slides were electrophoresed for 25 minutes at 300 mA (∼22-26 V). Slides were then incubated with neutralization buffer (0.4M Tris Base, pH 7.55) for 10 minutes at room temperature. Slides were fixed in 95% ethanol for 10 minutes and allowed to dry at room temperature. Cells were then stained with SYBR gold (Invitrogen), and images were captured with a Zeiss LSM 780 confocal microscope using a 63× lens. ∼50-100 cells were analyzed per slide using the OpenComet plug-in for Fiji 2. DNA damage was measured using the olive moment, which is the percentage of DNA within the tail multiplied by the length of the tail of the CAY10566-treated samples compared to the untreated samples [15].

### 2.11 Live/Dead assays in 3D cultures

Assays were performed on live ARCaP(M), PC3, 22Rv1, and C42B cells using Molecular Probes™ Live/Dead Viability/Cytotoxicity Kit (Invitrogen). Three-dimensional (3D) co-cultures of tumor cells were established as we described previously [16, 7] and cultured alone or on Transwell inserts and treated with either CAY10455 (1μM). EVO (50nM) or vehicle (0.1% DMSO) for 72 hours, re-treated and cultured for an additional 48 hours. For SCD knockdown experiments, spheroids were treated with scrambled siRNA or SCD siRNA for a minimum of 6 hours prior to combining with adipocyte cultures. For all experiments, coverslips were stained with 2µM Calcein AM and 5µM Ethidium homodimer-1 (Live/Dead Viability/Cytotoxicity Kit) for 30 minutes at room temperature, placed in PBS and immediately imaged by capturing z-stacks through the depth of structures using a Zeiss LSM 780 confocal microscope with a 40× water immersion objective. Live cells (green; Calcein AM) were captured using excitation at 488 nm and emission at 507 nm. Dead cells (red; Ethidium homodimer-1) were recorded using excitation at 488 nm and emission at 730 nm. 3D reconstruction and the sum of channel intensity were quantified using Volocity Software (Perkin Elmer, Waltham, MA). For each spheroid, the volume of live signal was obtained and shown as percent control of untreated cells.

### 2.12 RNAseq and pathway analyses

3’ RNA-seq (QuantSeq 3′ mRNA) was performed at the Wayne State University Applied Genomics Technology Center. RNA was collected from three biological replicates of PC3 and ARCaP(M) cells cultured alone or in Transwell with adipocytes, treated with 0.1% DMSO and 1μM CAY10566, as described above, and run on an Agilent TapeStation 2200 (Agilent Technologies, Santa Clara, CA) for quality control. Lexogen’s QuantSeq 3’mRNA-seq Library Prep Kit (FWD for Illumina) was utilized to build RNAseq libraries. The barcoded libraries were multiplexed at equimolar concentrations and sequenced with 50 bp single reads on an Illumina HiSeq-2500 run in rapid mode. Data were demultiplexed using Illumina’s CASAVA 1.8.2 software and reads were aligned to the human genomes [17]. Bioconductor package ‘edgeR’ was used to determine the differential gene expression during SCD inhibition in R [18]. ‘Enhanced Volcano’ package was used to create volcano plots representing the differentially expressed genes. LogFC cutoff was determined to be 1.5, and the p-value cutoff was determined to be 0.05 to show the significant differentially expressed genes (https://github.com/kevinblighe/EnhancedVolcano). Significant differentially expressed genes (p < 0.02 and logFC > 1) were analyzed using the “enrichGO” function in the ‘cluster profiler’ package in R to identify enriched pathways after FDR control using the MSigDB biological process (BP) gene set [19]. The top 20 BP pathways were shown on the dotplot after using ‘ggplot’ in R.

### 2.13 Intratibial injections and SCD inhibitor administration via oral gavage

Mice (minimum N=5/group) were placed on a high-fat diet (HFD; D12492i, Research Diets, New Brunswick, NJ) or low-fat diet (LFD; D12450Ji, Research Diets, New Brunswick, NJ) for 8 weeks. Mice were then intratibially injected with 5 × 10^5^ of ARCaP(M) or PC3 cells in PBS (20 µl, right tibia) or PBS alone (control, 20 µl, left tibia) under isoflurane inhalation anesthesia according to previously published procedures [8] and maintained on the diets for the duration of the study. For SCD inhibition, stock solutions of CAY10566 (TargetMol, Boston, MA) were prepared in sterile DMSO at 10mg/ml. Mice with confirmed ARCaP(M) or PC3 bone tumors (N=5/group) were administered CAY10566 (in 95% corn oil, 5% DMSO) at 5mg/kg via oral gavage daily. The experiment was repeated two more times. At the end of each study, mice were euthanized by CO_2_ inhalation, and control and tumor-bearing tibiae samples were fixed, decalcified, and embedded in paraffin.

### 2.14 Immunohistochemistry

Longitudinal sections (5μm) thick from the DMSO-treated and CAY10566-treated tumor-bearing tibiae were deparaffinized and examined by immunohistochemistry for expression and localization of ATF4 (1:200), ASNS (1:400), and γ-H2AX (1:200). Human metastatic bone prostate cancer samples from de-identified patients were obtained from the Biobanking Core at Karmanos Cancer Institute and sections examined for expression and localization for SCD (1:200, Bioss). ImmPRESS Anti-Rabbit Peroxidase Polymer Detection systems along with a NovaRED kit (Vector Labs, Newark, CA) as a substrate were used for the peroxidase-mediated immunostaining reaction.

### 2.15 Statistical analyses

For all analyses, data were presented as a mean of at least 3 experiments +SD and statistically analyzed using an unpaired student T-test.

### 2.16 Data Availability

RNAseq data will be deposited into NCBI Gene Expression Omnibus (GEO) repository (in process).

## 3. RESULTS

### 3.1 SCD expression varies across PCa cell lines and correlates with the effects of SCD inhibition on PCa growth and survival in 3D cultures

We have reported that SCD and its transcriptional regulators SREBP1 and SREBP2, along with several other genes in the desaturase pathway, are upregulated in patients with metastatic PCa and multiple myeloma, [4] two cancers highly regulated by marrow adiposity [4, 20, 21]. Based on this, we sought to determine whether SCD is specifically present in metastatic tumor cells and whether its levels are modulated by exposure to adipocytes. Western blot analysis of ARCaP(M), 22Rv1, PC3, and C42B cells grown in alone conditions revealed that ARCaP(M) and 22Rv1 cells **(Fig. 1A)** exhibit higher levels of SCD expression compared to PC3 and C42B cells **(Fig. 1B)**. Interestingly, SCD levels are further increased in ARCaP(M) and 22Rv1 cells grown in Transwell co-culture with adipocytes **(Fig. 1A),** whereas protein expression of SCD in PC3 and C42B cells appears to be reduced following adipocyte exposure (**Fig. 1B**). Immunohistochemical analyses of bone metastatic lesions from PCa patients confirm that SCD is indeed readily expressed by the majority of the metastatic tumor cells **(Supplementary** Fig. 1A). Notably, the level of SCD expression varies across patients, as corroborated by TaqMan RT-PCR analyses of bone metastatic cores **(Supplementary** Fig. 1B).

**Figure 1.**
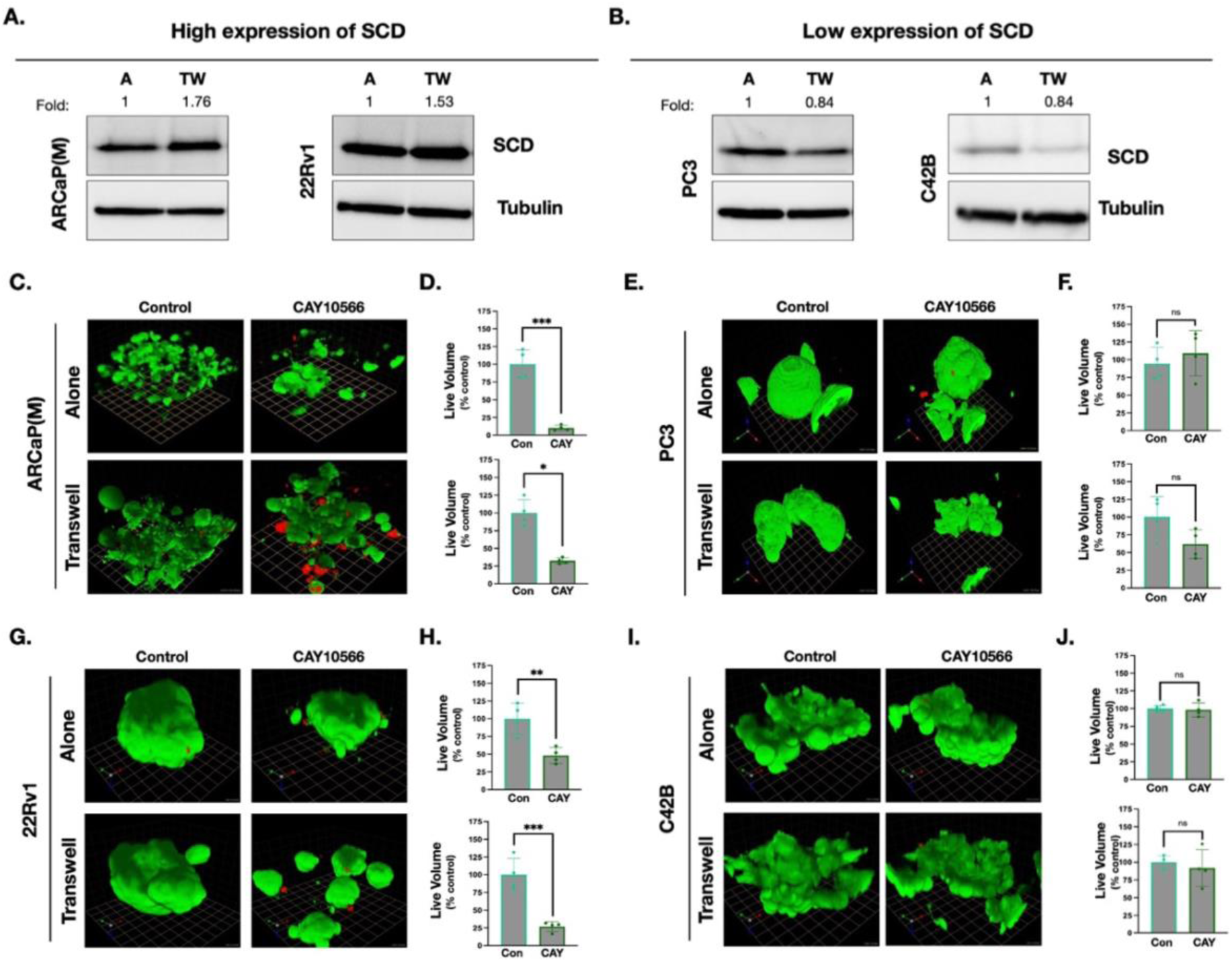
SCD expression levels vary between PCa cell lines and correlate with response to SCD inhibition in 3D culture. **A:** Western blot analyses of SCD expression in ARCaP(M), 22Rv1, **(B)** PC3, and C42B cells grown alone or in Transwell co-culture with adipocytes (TW). Fold change is calculated by comparing the expression levels of SCD to PCa cells in alone culture. **C, D:** ARCaP(M), PC3 **(E, F)**, 22Rv1 **(G, H)** and C42B **(I, J)** cells were grown in 3-dimensional cultures on reconstituted basement membrane (rBM) with a 2% Cultrex overlay. Live/Dead assay on 3D cultures grown alone **(top panels)** or with adipocytes **(bottom panels)** and treated with vehicle control (0.1% DMSO) or 1μM CAY10566. Live cells (green fluorescence; calcein AM); dead cells (red fluorescence; ethidium homodimer). Quantification of live (green) spheroid volume compared to control for ARCaP(M) **(D)**, PC3 **(F)**, 22Rv1 **(H)** and C42B **(J)** cells grown alone **(top panel)** or with adipocytes **(bottom panel)**; * p < 0.05, ** p < 0.01, *** p < 0.001 and ns = not significant.

To determine if the level of SCD expression directly impacts sensitivity to SCD inhibition, we cultured ARCaP(M), 22Rv1, PC3, and C42B 3D spheroids alone or with adipocytes in the absence or presence of SCD-selective inhibitor CAY10566 (1μM) and performed the Live/Dead assay **(Fig. 1C-J)**. A significant decrease in live volume (calcein AM; green fluorescence) was observed in ARCaP(M) **(Fig. 1C, D top panels)** and 22Rv1 spheroids **(Fig. 1G, H top panels)** cultured alone in the presence of SCD inhibitor. Notably, even when grown in Transwell co-culture with adipocytes, which we have shown to diminish response to therapies, such as docetaxel [16, 7], ARCaP(M), and 22Rv1 spheroids, remained sensitive to SCD inhibition **(Fig. 1C, D bottom panels and G, H bottom panels)**. In contrast to pharmacological SCD inhibition, siRNA-mediated SCD knockdown (verified in **Supplementary** Fig. 2) reduced the volume of ARCaP(M) spheroids cultured alone, but had no significant impact on the spheroid size when cultured in the presence of adipocytes **(Supplementary** Fig. 3A, B). Since SCD is also expressed by adipocytes and is essential for *de novo* lipogenesis [22, 23], our results suggest that inhibition of SCD in both the tumor cells and the adipocytes might be needed for the robust tumor response. Interestingly, in contrast to ARCaP(M) spheroids, PC3 and C42B 3D spheroids cultured in alone conditions **(Fig. 1E, F, I, J top panels)** or in the presence of adipocytes **(Fig. 1E, F, I, J bottom panels)** showed no significant change in live volume following the pharmacological inhibition with CAY10566. No change in live volume was also demonstrated in PC3 spheroids cultured alone or in the presence of adipocytes during siRNA-mediated SCD knockdown **(Supplementary** Fig. 3C, D).

### 3.2 Prostate cancer cells control lipid peroxidation induced by bone marrow adipocytes through SCD activity

Cancer cells often rewire lipid metabolism to fuel their proliferation and adapt to stress [24, 25], producing various lipid species that serve as substrates for lipid peroxidation (LPO) [26]. We have previously determined that PCa cells stimulate lipid hydrolysis in adipocytes and subsequently utilize the released lipids to fuel their growth and survival [7]. To determine if bone marrow adipocyte-derived lipids affect LPO levels in PCa SCD-amplified cells differently than non-amplified cells and, in turn, rely on SCD to adapt to LPO, we utilized the LPO sensor BODIPY C11. Lipid radical formation in ARCaP(M) and PC3 cells in alone and Transwell conditions with bone marrow adipocytes was measured based on the shift from red (561/590 nm) to green (488/510 nm) fluorescence. Our results showed significant induction in LPO levels indicated by an increase in green fluorescence occurring in both ARCaP(M) and PC3 cells when exposed to adipocytes for 24 hours compared to cells in monoculture **(Fig. 2A-D).** Intriguingly, a reduction in LPO levels was observed in ARCaP(M) **(Fig. 2A, B)** and PC3 **(Fig. 2C, D)** cells exposed to adipocytes for 48 hours, as compared to the 24-hour time point, suggesting the activation of a potential adaptive response in PCa cells to mitigate harmful lipid peroxides. Notably, the magnitude of the adaptive response differed between the two cell lines, with PC3 cells demonstrating a more significant drop in LPO levels at 48 hours, almost to the levels of PC3 cells in alone culture **(Fig. 2A-D)**.

**Figure 2.**
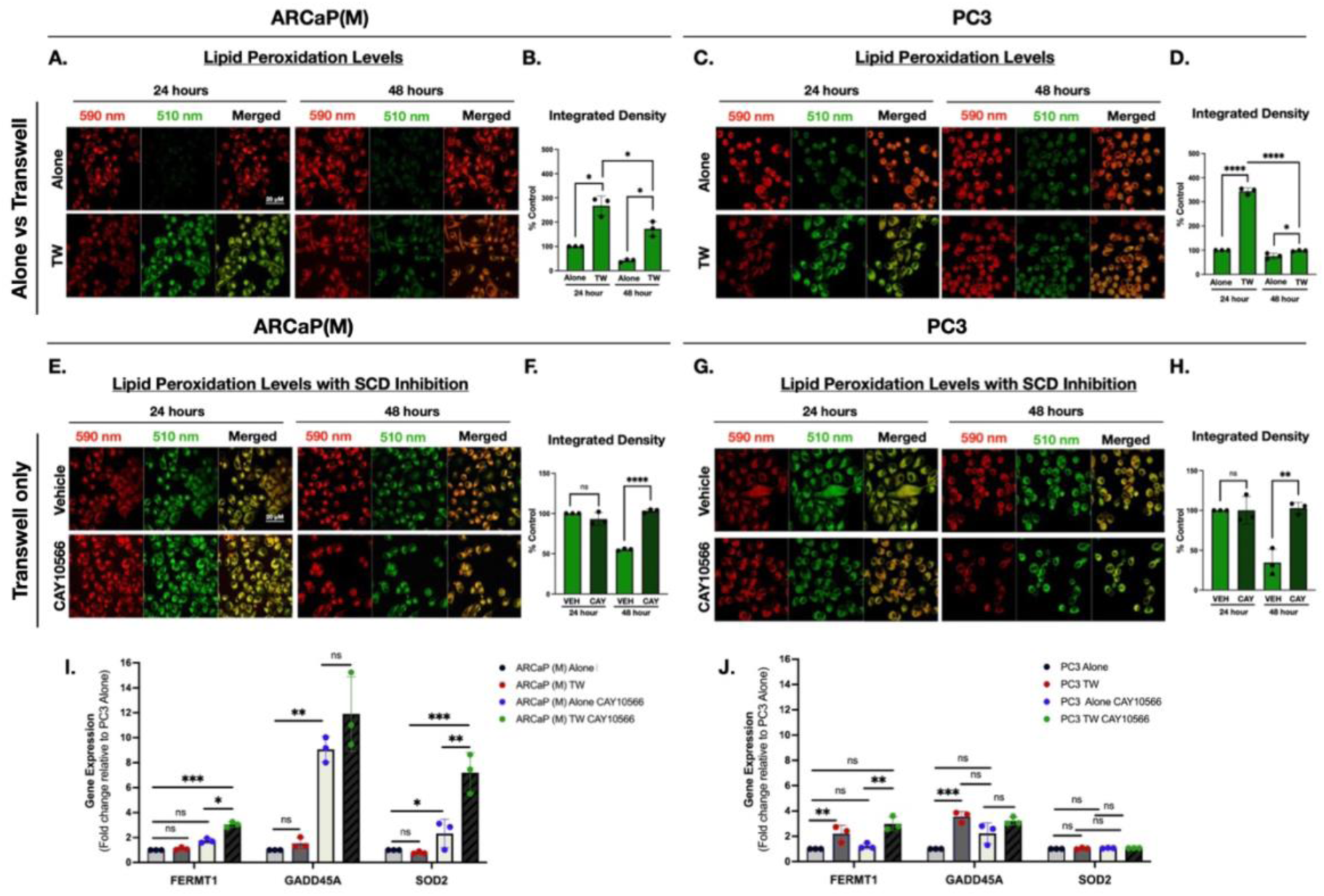
PCa cells exhibit reduced lipid peroxidation (LPO) levels during prolonged adipocyte exposure and increased LPO levels by SCD inhibition. ARCaP(M) **(A, B)** and PC3 **(C, D)** cells were grown alone or in TW co-culture with adipocytes for 24 and 48 hours. BODIPY C-11 staining was performed to examine LPO levels. An increase in LPO is indicated by a shift of fluorescence emission peak from ∼590 nm (red) towards ∼510 nm (green); 40× images. Quantification of ∼510 nm shift for ARCaP(M) **(B)** and PC3 cells **(D),** indicating reduced LPO at 48 hrs as compared to 24 hrs. ARCaP(M) **(E, F)** and PC3 **(G, H)** cells were grown in TW co-culture with adipocytes and treated with vehicle control (0.1% DMSO) or 1μM CAY10566 for 24 and 48 hours. **F:** Quantification of ∼510 nm shift for ARCaP(M) cells and PC3 cells **(H),** indicating an increase in LPO upon treatment with CAY10566 compared to vehicle control. **I, J:** TaqMan RT-PCR analysis for mRNA expression of *FERMT1, GADD45A* and *SOD2* in ARCaP(M) **(I)** and PC3 **(J)** cells grown in alone conditions or in TW co-culture with adipocytes and treated with vehicle control (0.1% DMSO) or 1 μM CAY10566. Data represent at least 3 experiments; *p < 0.05, ** p < 0.01, *** p < 0.001, **** p < 0.0001 and ns = not significant.

To examine if SCD may have a direct role in modulating LPO levels in PCa cells during adipocyte exposure, we pharmacologically inhibited SCD activity using CAY10566 and monitored changes in LPO using the BODIPY C11 sensor. The quantification of BODIPY C11 fluorescence following 24-hour exposure to CAY10566 revealed that LPO levels, which are already increased in ARCaP(M) and PC3 cells by 24-hour adipocyte exposure **(Fig. 2A-D)**, were not further affected by SCD inhibition **(Fig. 2E-H)**. However, treatment with CAY10566 for 48 hours not only prevented the adaptive decrease in peroxidation levels we typically observe at this time point, but it significantly increased the amount of green BODIPY C11 fluorescence, indicating augmented levels of lipid peroxides with prolonged SCD inhibition **(Fig. 2E-H)**. ARCaP(M) cells in monoculture exhibited no change in LPO levels during SCD inhibition by CAY10566 at 24 or 48 hours **(Supplementary** Fig. 4A-C). However, PC3 cells in monoculture had a slight increase in LPO levels at 24 hours but not at 48 hours when treated with CAY10566 **(Supplementary** Fig. 4D-F). siRNA-mediated knockdown of SCD in PCa cells exposed to adipocytes (verified in **Supplementary** Fig. 2) led to similar increases in BODIPY C11 fluorescence **(Supplementary** Fig. 5). Notably, PCa cells cultured in the absence of adipocytes exhibited minimal changes in LPO levels following either pharmacological inhibition or siRNA-mediated knockdown of SCD **(Supplementary** Fig. 6), underscoring the role of SCD in promoting an adaptive response within a lipid-rich environment.

Consistent with these findings, SCD inhibition led to an upregulation of genes encoding proteins that play a role in cellular defense mechanisms against damaging stress. Specifically, mRNA levels of *Superoxide Dismutase 2* (*SOD2*), *FERM Domain Containing Kindlin 1* (*FERMT1*), and *Growth Arrest and DNA Damage Inducible Alpha* (*GADD45A*) were significantly higher in ARCaP(M) cells treated with CAY10566, as compared to cells treated with vehicle control, both under monoculture and co-culture conditions **(Fig. 2I)**. Intriguingly, unlike in ARCaP(M) cells, *SOD2* levels in PC3 cells remained completely unaffected by either co-culture with adipocytes or SCD inhibition. Moreover, *FERMT1* and *GADD45A* levels increased in PC3 cells co-cultured in Transwell with adipocytes, but there were no additional changes in the expression of these stress-related genes during SCD inhibition **(Fig. 2J)**. 22Rv1 cells, which responded to SCD inhibition in 3D culture similarly to ARCaP(M) cells, showed increased *GADD45A* expression during SCD inhibition, mirroring the impact on this gene in ARCaP(M) cells **(Supplementary** Fig. 7A). This effect was absent in C42B cells **(Supplementary** Fig. 7B), which, like PC3 cells, exhibited limited sensitivity to SCD inhibition in 3D culture. Similar to PC3 cells, C42B cells showed increased *GADD45A* expression upon exposure to adipocytes, yet displayed minimal transcriptional changes in response to SCD inhibition. These findings suggest potential cell-line dependent differences in adaptability to adipocyte-mediated LPO, cellular stress, and the loss of SCD activity.

### 3.3 SCD inhibition induces distinct changes in the transcriptome of PCa cells

To investigate the molecular mechanisms responsible for the differential responses of PCa cell lines to SCD inhibition, we conducted RNAseq analyses on ARCaP(M) and PC3 cells cultured alone in the presence or absence of CAY10566. Our results showed that pharmacological SCD inhibition significantly altered the transcriptome of ARCaP(M) cells with a substantially larger number of differentially expressed genes (DEGs) identified (**Fig. 3A)** as compared to PC3 cells **(Fig. 3B)**, which we have shown to be unresponsive to SCD inhibition in 3D culture. Through gene ontology (GO) analysis, we determined that in ARCaP(M) cells under alone conditions, SCD inhibition predominantly affected pathways involved in the endoplasmic reticulum stress, the unfolded protein response, and apoptosis **(Fig. 3C)**. Furthermore, a significant enrichment for mTOR signaling was also observed, although it did not rank among the top 20 pathways displayed in the dotplot in **Fig. 3C**. In contrast, PC3 cells under the same conditions exhibited enrichment of pathways primarily involved in oxidative stress and hypoxia response following SCD inhibition **(Fig. 3D)**.

**Figure 3.**
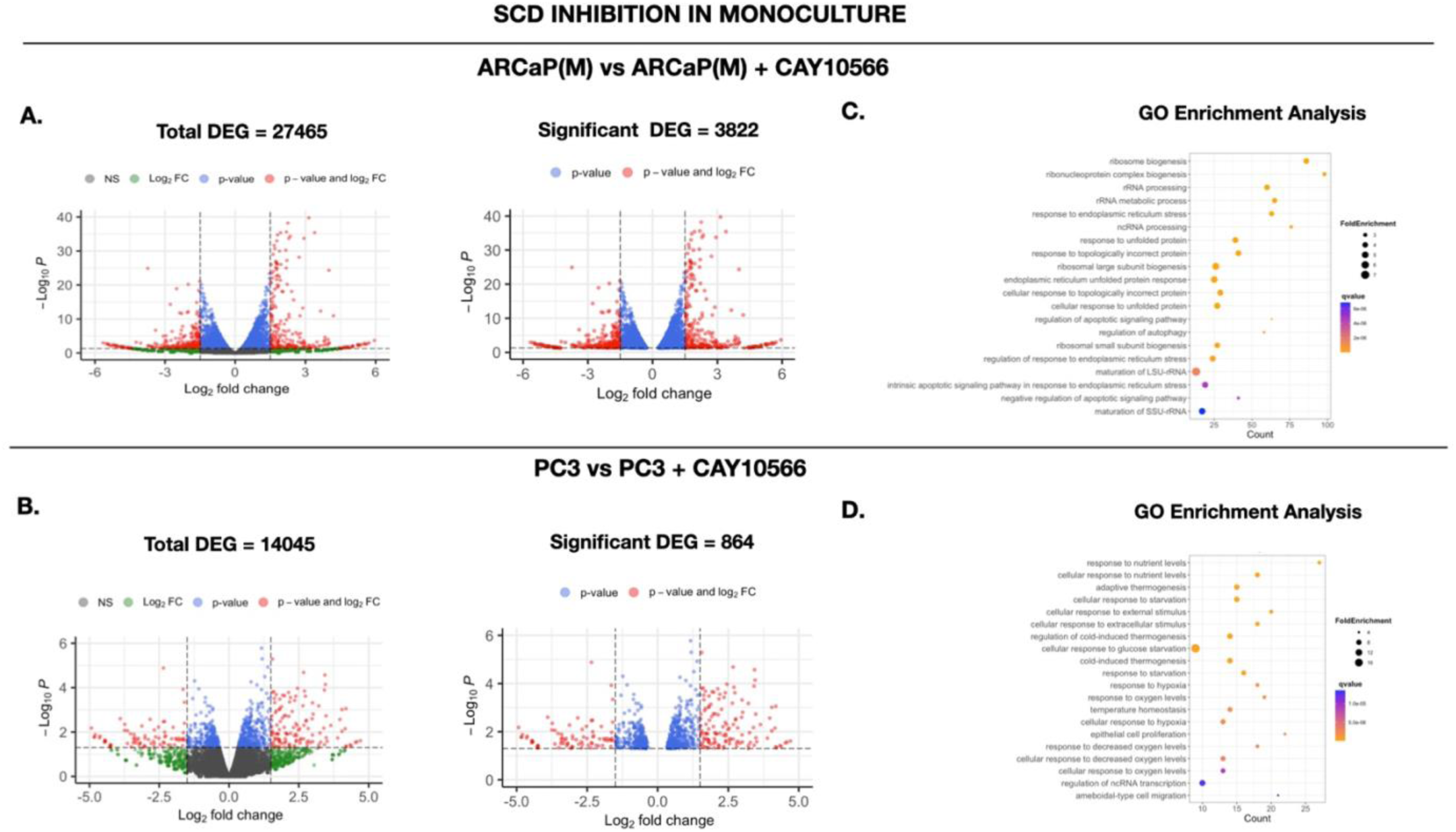
SCD inhibition of PCa cells in monoculture affects the transcriptome and induces pathway changes. **A:** EnhancedVolcano plot indicating the differentially expressed genes (DEG) (left) and significant DEG (right) in ARCaP(M) and **(B)** PC3 cells grown in alone conditions in the presence or absence of 1μM CAY10566. **C:** Gene ontology pathway enrichment analyses using enrichGO of significant DEGs (p < 0.02) in ARCaP(M) and PC3 **(D)** cells using the biological processes function. The count on the x-axis depicts the number of genes in each pathway, Fold Enrichment determines the dot size, and the gradient color indicates q-value; increasing darkness of color indicates lower q-value.

### 3.4 SCD inhibition alters distinct pathways in PCa cells exposed to adipocytes compared to PCa cells in monoculture

Given that adipocytes dampened the response of ARCaP(M) spheroids to SCD inhibition, we next investigated how SCD inhibition alters the transcriptome of PCa cells in Transwell co-culture with adipocytes. Our data show that ARCaP(M) cells treated with CAY10566 in the presence of adipocytes exhibited a significant number of DEGs compared to ARCaP(M) cells treated with CAY10566 alone, suggesting that distinct genes and pathways may be affected in Transwell co-culture conditions **(Fig. 4A)**. We demonstrated this by GO analysis, which revealed that ARCaP(M) cells treated with CAY10566 in the presence of adipocytes show enrichement in pathways related to DNA damage, cell cycle and the unfolded protein response, compared to ARCaP(M) cells treated with CAY10566 alone **(Fig. 4B)**. In contrast, PC3 cells in Transwell co-culture with adipocytes displayed only a small number of DEGs relative to PC3 cells in monoculture **(Fig. 4C)**. GO analyses of PC3 cells indicated low pathway enrichment, primarily involving the ER stress response **(Fig. 4D)**. These findings suggest that SCD inhibition alters the transcriptome of PCa cells differently depending on the presence of marrow adipocytes, with a more pronounced effect in SCD-amplified cell lines like ARCaP(M).

**Figure 4.**
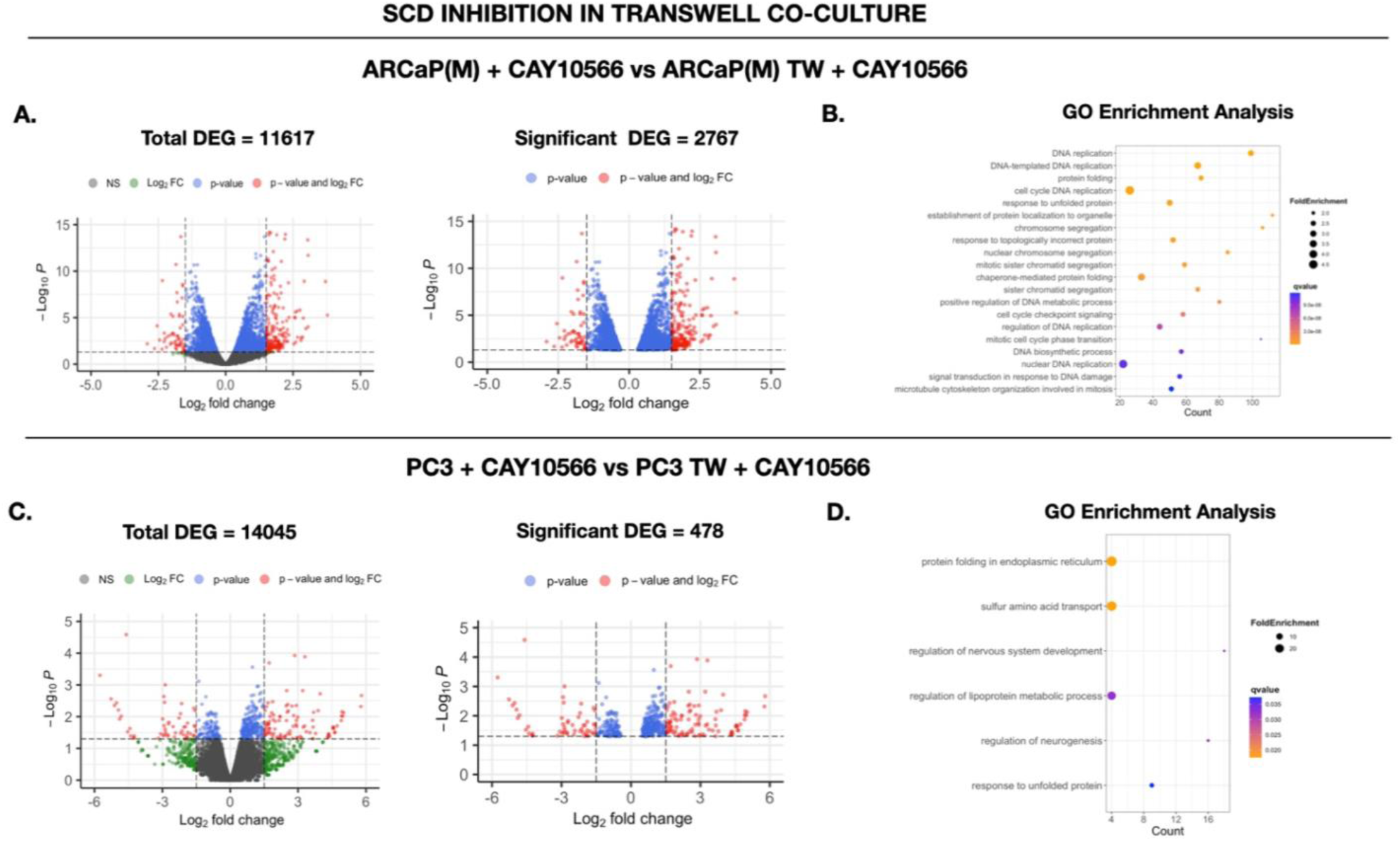
SCD inhibition in PCa cells exposed to adipocytes induces different transcriptomic changes and enriches distinct pathways compared to PCa cells in monoculture with SCD inhibition. **A:** EnhancedVolcano plot indicating the differentially expressed genes (DEG) (left) and significantly expressed DEG (right) in ARCaP(M) and **(B)** PC3 cells treated with 1μM CAY10566. **C:** Gene ontology pathway enrichment analyses using enrichGO of significant DEGs (p < 0.02) in ARCaP(M) and PC3 cells **(D)** treated with 1μM CAY10566 in either alone conditions or in Transwell co-culture with adipocytes using the biological processes function. The count on the x-axis depicts the number of genes in each pathway, Fold Enrichment determines the dot size, and the gradient color indicates q-value; increasing darkness of color indicates lower q-value.

### 3.5 Differential impact of SCD inhibition on ER stress and mTOR signaling in sensitive and non-sensitive cell lines

Since our RNAseq analyses exposed ER stress signaling as a highly enriched pathway in ARCaP(M) cells in alone and Transwell conditions following SCD inhibition, we examined protein expression of a major ER stress regulator, activating transcription factor 4 (ATF4), in ARCaP(M) and PC3 cells grown in alone conditions and Transwell co-culture with adipocytes in the presence or absence of SCD inhibitor **(Fig. 5A, C)** or siRNA targeting SCD **(Supplementary** Fig. 8A, B). Our results revealed major differences in the effects of SCD inhibition on ATF4 expression between the two cell lines. Specifically, consistent with our previously reported results [11], Transwell co-culture with adipocytes led to a significantly greater increase in ATF4 levels in PC3 cells compared to ARCaP(M) cells **(Fig. 5A, C)**. Notably, in PC3 cells, treatment with CAY10566 or siRNA-mediated SCD knockdown further elevated ATF4 levels in the presence of adipocytes, with minimal impact under monoculture conditions. In contrast, SCD inhibition in ARCaP(M) cells resulted in highly augmented ATF4 levels regardless of the presence or absence of adipocytes **(Fig. 5A, C** and **Supplementary** Fig. 8A, B). These notable increases in ATF4 levels, especially in ARCaP(M) cells, were accompanied by a decrease in the protein expression of binding immunoglobulin protein (BIP), indicative of active unfolded protein response signaling [27, 28]. Interestingly, 22Rv1 cells, which displayed sensitivity to SCD inhibition in 3D culture similar to that of ARCaP(M) cells (**Fig. 1**), also showed comparable changes in ATF4 and BIP protein levels upon SCD inhibition or siRNA-mediated knockdown. **(Supplementary** Fig. 8C, D). In contrast, the effects on ATF4 and BIP levels in C42B cells resembled those observed in PC3 cells **(Supplementary** Fig. 8E, F). These findings highlight the potential importance of ER stress response in mediating sensitivity to SCD inhibition.

**Figure 5.**
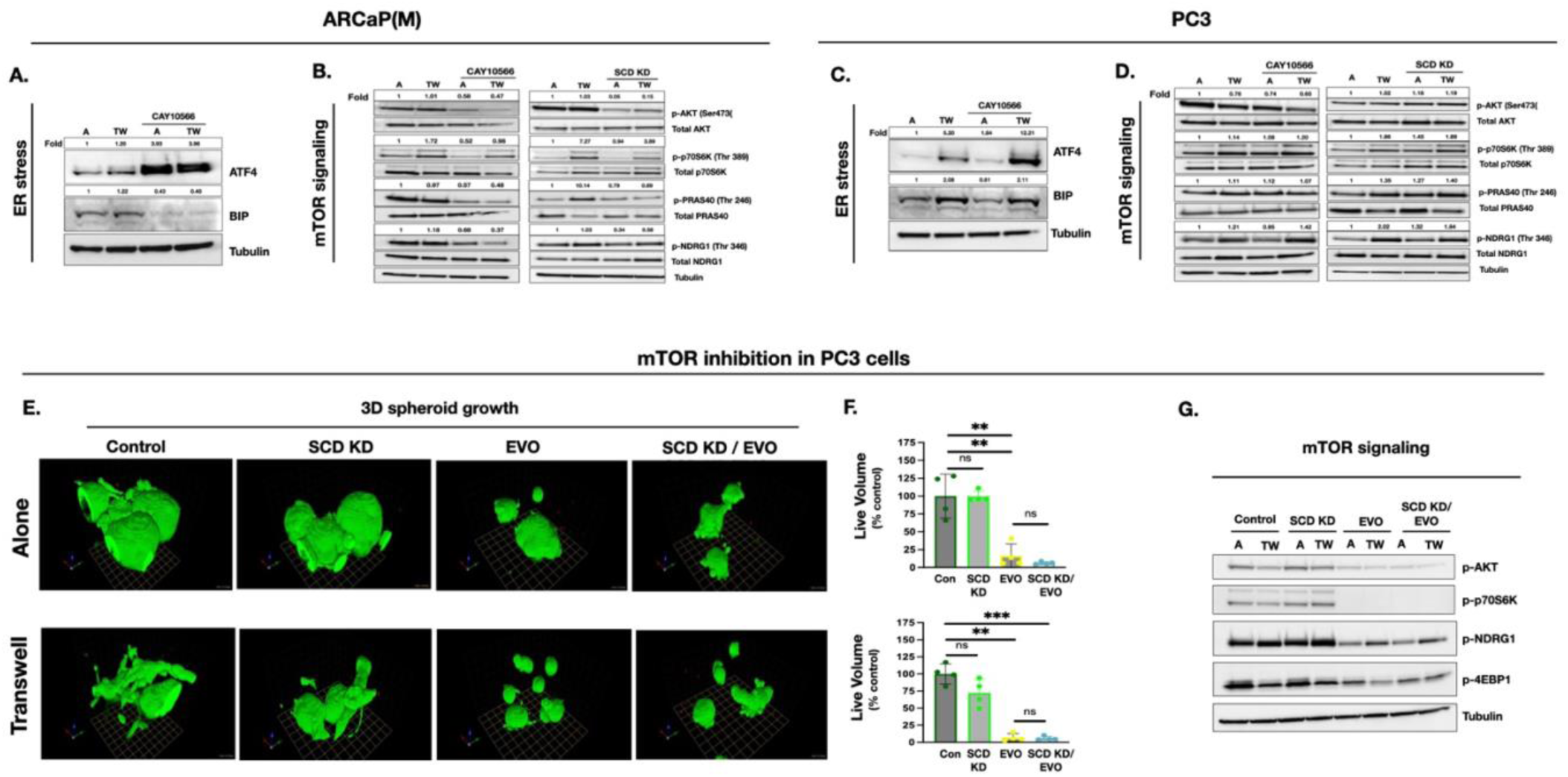
SCD loss increases ER stress levels and decreases mTOR signaling in only ARCaP(M) cells and PC3 cells cannot be sensitized to SCD inhibition by inhibition of mTOR signaling. **A:** Immunoblot analyses of ER stress markers ATF4 and BIP in ARCaP(M) and PC3 cells **(C)** grown alone or in TW co-culture with adipocytes, treated with 1μM CAY10566. ARCaP(M) **(B)** and PC3 **(D)** cells were cells grown alone or TW co-culture with adipocytes and treated with vehicle control (0.1% DMSO) or 1μM CAY10566 and scrambled siRNA or siRNA targeting SCD for 48 hours and subjected to immunoblot analysis of total and phosphorylated downstream mTOR proteins: AKT, P70S6K, PRAS40 and NDRG1. **E:** PC3 cells were grown in 3-dimensional cultures on reconstituted basement membrane (rBM) with a 2% Cultrex overlay. Live/Dead assay on 3D cultures grown alone **(top panels)** or with adipocytes **(bottom panels)** and treated with scrambled siRNA and vehicle control (0.1% DMSO) (Control), siRNA targeting SCD and 50nM Everolimus (EVO). Live cells (green fluorescence; calcein AM); dead cells (red fluorescence; ethidium homodimer. **F:** Quantification of live (green) spheroid volume compared to control for PC3 cells grown in monoculture **(top panel)** or with adipocytes **(bottom panel). G:** PC3 cells were grown in alone conditions or in TW co-culture with adipocytes and treated with vehicle control (0.1% DMSO) or 10nM EVO for 48 hours and subjected to immunoblot analysis of phosphorylated downstream mTOR proteins: AKT, P70S6K, NDRG1, and 4EBP1. Data represent at least 3 experiments; ** p < 0.01; *** p < 0.001 and ns = not significant.

In addition to a strong ER stress response, gene ontology enrichment analysis of ARCaP(M) cells exposed to SCD inhibition revealed a significant enrichment of the mTOR signaling pathway, which is crucial for cancer cell growth and survival [29]. Specifically, we observed a visible reduction in phosphorylation of downstream mTOR proteins AKT, p70S6K, PRAS40, and NDRG1 in ARCaP(M) cells treated with SCD inhibitor CAY10566 and with siRNA targeting SCD **(Fig. 5B)**. This was also seen in ARCaP(M) cells with the use of another SCD inhibitor A-939572 **(Supplementary** Fig. 9A). In contrast to ARCaP(M) cells, there was no observable impact on downstream mTOR proteins in PC3 cells treated with SCD inhibitor CAY10566, SCD siRNA **(Fig. 5D)** or SCD inhibitor A-939572 **(Suppmentary Fig. 9B)**, a result that aligns with the limited effect of SCD inhibition on PC3 viability in 3D assays. We have shown previously that adipocyte exposure can increase expression of lipid transporters such as fatty acid binding protein 4 (FABP4) and cluster of differentiation 36 (CD36) in PCa cells [30]. Given that lipids and the upregulation of lipid transporters can modulate cell growth and survival via the mTOR pathway [31-33], we used BODIPY 493/503 dye to assess whether lipid droplet accumulation or lipid transporter expression could account for differences in viability and mTOR signaling between ARCaP(M) and PC3 lines. Upon exposure to adipocytes, PC3 cells accumulated significantly more lipid droplets than ARCaP(M) cells **(Supplementary** Fig. 10A, B). Additionally, Transwell co-culture induced the expression of *CD36* and *FABP4* in PC3 but not ARCaP(M) cells **(Supplementary** Fig. 10C, D), and this expression was further increased with SCD inhibition in PC3 cells, with no significant change observed in ARCaP(M) cells **(Supplementary** Fig. 10C, D). These findings suggest that increased lipid uptake and transporter expression in PC3 cells may help sustain mTOR activity and cell viability, thereby reducing sensitivity to SCD inhibition.

To determine whether inhibiting mTOR signaling could sensitize PC3 cells to SCD inhibition, we performed Live/Dead assay on PC3 spheroids treated with either scrambled control siRNA or SCD siRNA, in the presence or absence of mTORC1 inhibitor, Everolimus (EVO). Our results showed that PC3 cells are highly sensitive to mTOR inhibition alone and that the combination treatment with SCD inhibition does not provide additional benefit (**Fig. 5E, F**). Immunoblot analysis revealed a strong reduction in the levels of pAKT and mTOR proteins following treatment with EVO, regardless of SCD knockdown (**Fig. 5G**), a result consistent with constitutive activation and dependence on mTOR signaling in PC3 cells [34].

### 3.6 Inhibition of SCD activity increases the expression of DNA repair genes and induces DNA damage

Given the known links between lipid-driven metabolic stress, ER stress, and genomic instability, we next investigated whether SCD inhibition might elicit DNA damage response in CAY10566-sensitive PCa cells [35-37]. Mining of our RNA sequencing data from ARCaP(M) cells exposed to adipocytes in the presence of CAY10566 revealed a striking upregulation of genes involved in DNA repair and the cell cycle (**Fig. 4B)**. Intriguingly, this transcriptional response was not observed in PC3 cells under the same conditions **(Fig. 4D)**. To determine if the transcriptome changes in ARCaP(M) cells following SCD inhibition were a direct consequence of DNA damage, we performed immunofluorescence staining for γ-H2AX, a marker of DNA double-stranded breaks. SCD inhibition by CAY10566 significantly increased γ-H2AX fluorescence intensity in ARCaP(M) **(Fig. 6A)** and 22Rv1 **(Fig. 6B)** cells exposed to adipocytes with no detectable increase in PC3 **(Fig. 6C)** and C42B cells **(Fig. 6D).** Interestingly, ARCaP(M) cells exposed to adipocytes in the absence of SCD inhibition exhibited fewer γ−H2AX foci compared to cells in monoculture **(Fig. 6A)**, highlighting a protective effect from adipocyte interaction. This observation was further supported by an alkaline comet assay, which detects both single- and double-stranded DNA breaks. Specifically, ARCaP(M) cells co-cultured with adipocytes and treated with vehicle control displayed fewer comet tails, indicative of reduced DNA damage, compared to ARCaP(M) cells cultured in alone conditions **(Fig. 6E)**. ARCaP(M) cells subjected to SCD inhibition via treatment with CAY10566 showed a significant increase in DNA strand breaks, as evidenced by a higher olive moment in the comet assay **(Fig. 6E, F)**.

**Figure 6.**
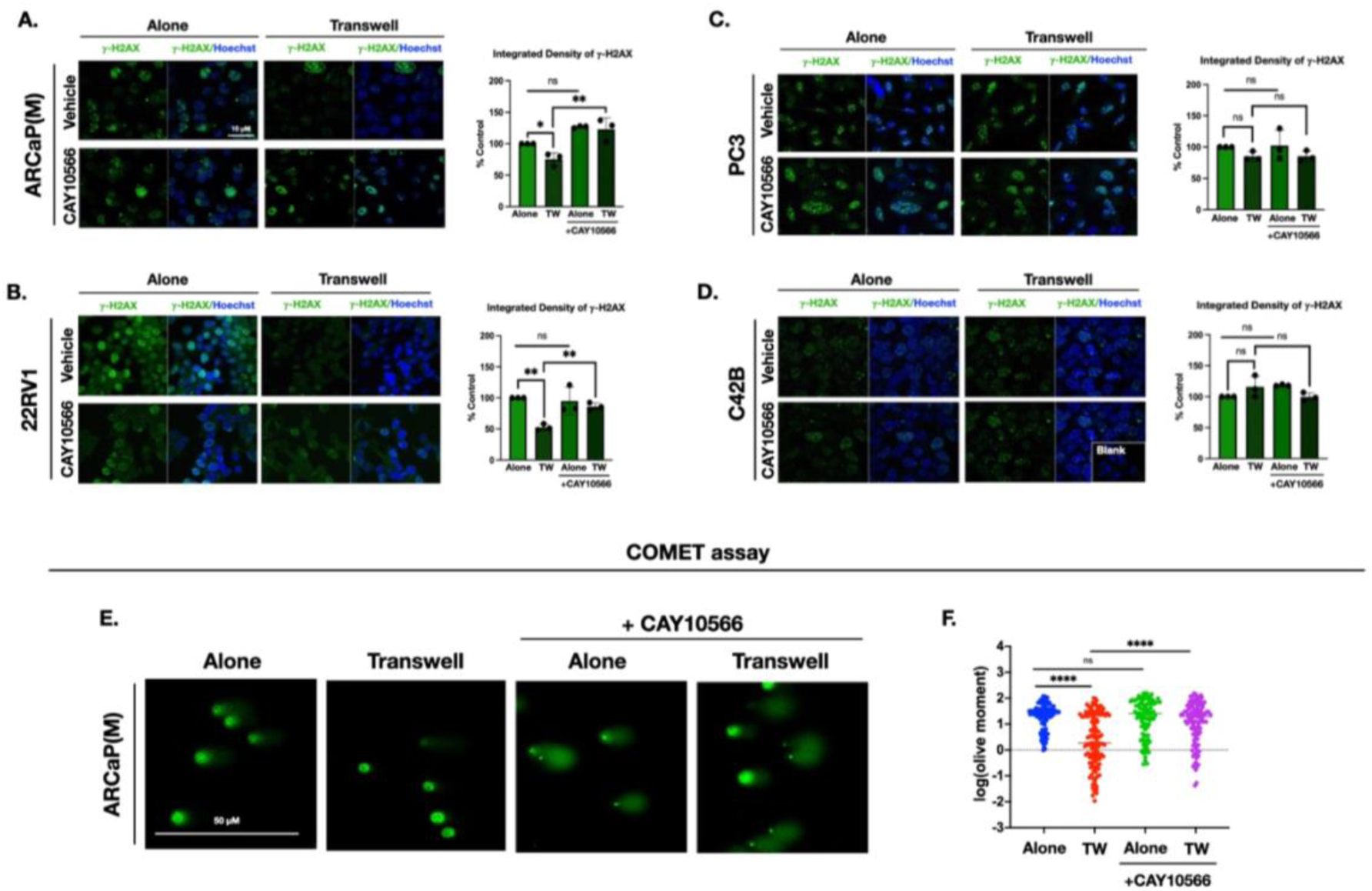
SCD inhibition induces DNA damage in ARCaP(M) and 22Rv1 cells but does not significantly affect DNA damage response in PC3 and C42B cells. **A-D:** Immunofluorescence staining (green) of γ-H2AX in ARCaP(M) **(A)**, 22Rv1 **(B)**, PC3 **(C)** and C42B **(D)** cells grown alone or in TW co-culture with adipocytes and treated with vehicle control (0.1% DMSO) or 1μM CAY10566 for 48 hours; 63× images; Hoechst dye (blue) was used to stain the nuclei. Quantification of γ-H2AX fluorescence in ARCaP(M) **(A)**, 22Rv1 **(B)**, PC3 **(C)** and C42B **(D)** cells; data represent at least 3 experiments; **E:** ARCaP(M) cells grown alone or in TW co-culture were treated with vehicle control (0.1% DMSO) or 1μM CAY10566 and subjected to modified alkaline comet assay and stained with SYBR gold; 20× images. **F:** DNA damage was quantified for all conditions using the olive moment for ∼50-100 cells for each condition in the OpenComet plug-in for Fiji 2. Data was log-transformed to normalize the distribution; * p < 0.05; ** p < 0.01; **** p < 0.0001 and ns = not significant.

### 3.7 SCD inhibition suppresses growth and induces ER stress and DNA damage in prostate bone tumors

To examine the *in vivo* response to SCD inhibition in a lipid-rich environment, we utilized our well-established intratibial model of tumor growth and induced marrow adiposity through HFD feeding [14, 7, 11, 12, 8]. Mice with established intratibial ARCaP(M) or PC3 tumors (three weeks post-injection) were treated with the SCD inhibitor (CAY10566, 5mg/kg) or vehicle (5% DMSO) via oral gavage for four weeks. Histological analyses of resulting bone tumor sections revealed a significant decrease in ARCaP(M) bone tumor area in CAY10566-treated mice compared to vehicle controls **(Fig. 7A, B)**. In contrast, no significant effect of SCD inhibition on PC3 bone tumor area was observed **(Fig. 7C, D),** consistent with the *in vitro* findings of the insensitivity of PC3 cells to SCD inhibition. Notably, an increase in γ-H2AX foci staining was observed in ARCaP(M) but not PC3 bone tumors following CAY10566 treatment **(Fig.7E, F)**.

**Figure 7.**
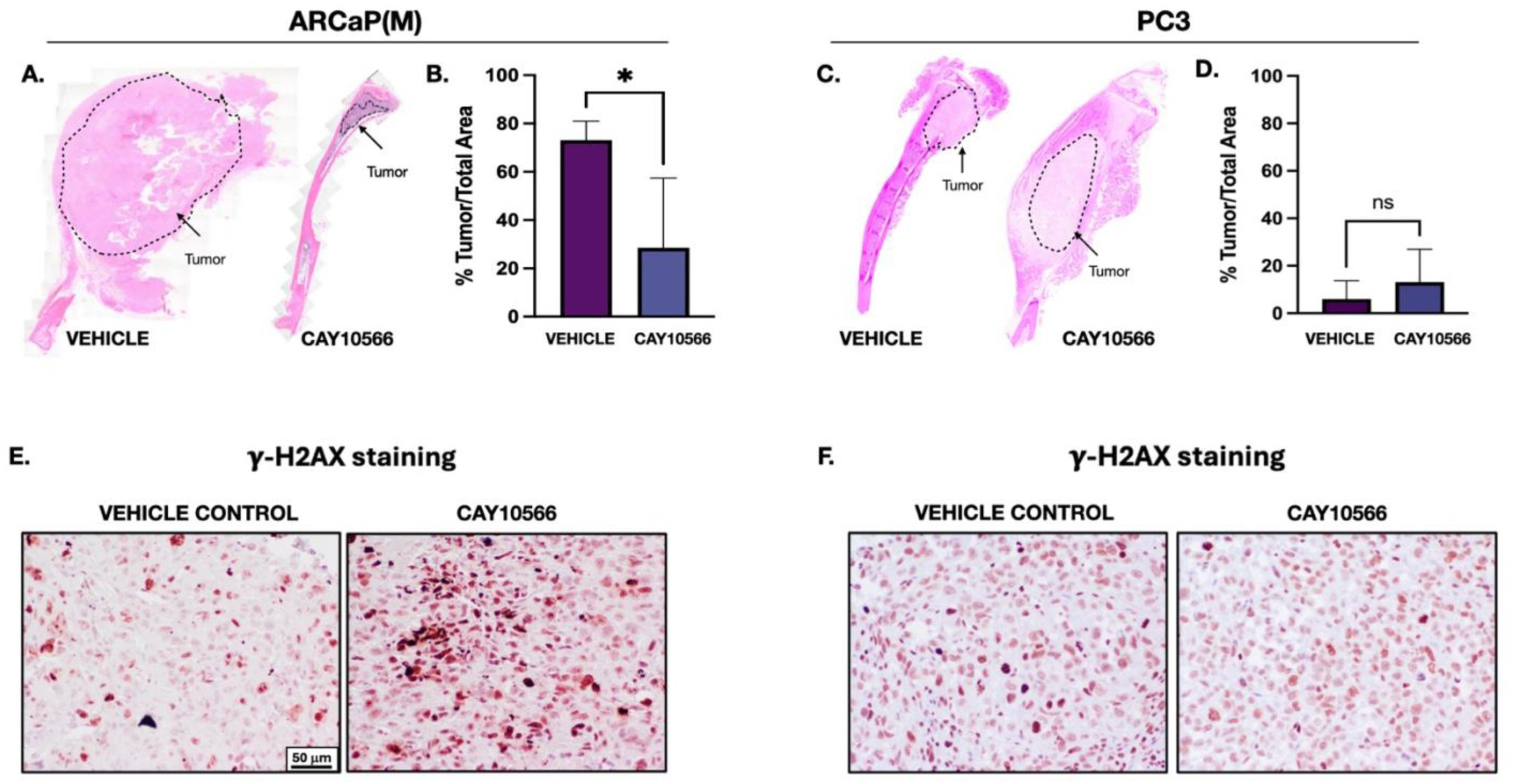
SCD inhibition *in vivo* reduces ARCaP(M) tumor size. A,C: Histological examination of ARCaP(M) **(A)** and PC3 **(C)** tumors from mice treated with vehicle (5% DMSO) (left) or 5 mg/kg CAY10566 (right) by hematoxylin and eosin (H&E) staining; residual tumors in responder mice are indicated by the dotted line. **B, D:** Quantification of percent tumor area for vehicle (VEH) and CAY10566-treated ARCaP(M) **(B)** and PC3 **(D)** tumors from mice. **E, F:** Immunohistochemical (NovaRed) staining of and γ-H2AX protein in ARCaP(M) **(E)** and PC3 **(F)** bone tumors from vehicle-treated (left) and CAY10556-treated (right) mice; 20× images, * p < 0.05, ns = not significant.

Additionally, ARCaP(M) tumors from treated mice showed elevated expression of ER stress marker ATF4 and its downstream target asparagine synthetase (ASNS), compared to vehicle-treated controls **(Supplementary** Fig. 11A-D). Together, these results support our *in vitro* findings and highlight the role of SCD in PCa survival and progression in bone.

## 4. DISCUSSION

There is an unmet clinical need to identify molecular drug targets for metastatic prostate cancer in bone, primarily due to the limited understanding of the mechanisms that promote tumor growth and survival within the bone microenvironment. The aim of the present study was to gain a deeper understanding of the adaptive mechanisms that drive metastatic progression in the lipid-rich bone marrow niche. For the first time, we show that tumor cells adapt to the harsh bone microenvironment and overcome lipid peroxidation induced by bone marrow adipocytes by upregulating the lipid desaturase SCD. We also uncovered that dependence on SCD activity to mitigate lipid peroxidation in the harsh bone microenvironment varies across PCa cell lines, and that cell lines with higher SCD dependence may be more sensitive to DNA-damaging agents. In PCa cell lines that respond to SCD inhibition, we observe reduced mTOR signaling and transcriptomic alterations in DNA repair pathways, consistent with DNA damage induced by SCD deficiency. We also demonstrate that SCD inhibition significantly reduces the size of experimental bone tumors in mice, supporting its potential as a therapeutic strategy for patients with bone metastatic PCa. This is the first report to present evidence that SCD is a tumor-promoting enzyme in bone that regulates lipid peroxidation levels in the hostile bone microenvironment.

An increasing number of studies highlight the role of LPO in promoting tumor progression [38]. SCD has been implicated in tumor cell evasion from lipid peroxide-induced cell death, specifically by protecting cells against ferroptosis [39, 40]. Lipid peroxides are thought to arise from ROS species targeting PUFAs [41]. Notably, MUFAs are believed to be key in safeguarding PUFAs from peroxidation [42]; however, SCD’s direct role in this process has yet to be established. Here, our findings suggest that expression of SCD in PCa cell lines may help mitigate adipocyte-induced LPO levels, consistent with our observation that SCD inhibition leads to elevated LPO levels. We also found that increased LPO levels robustly induce ER stress in PCa cell lines exposed to adipocytes, and this effect is more dramatic in SCD-insensitive cell lines such as PC3 or C42B. These cell lines, which also exhibit higher expression of oxidative stress genes upon adipocyte exposure, appear to mount a stronger adaptive response to fat cells and this preemptive adaptation may render them more resistant to SCD inhibition. In contrast, SCD-sensitive lines show minimal induction of ER stress proteins and oxidative stress genes under adipocyte exposure, but drastically increase these genes during SCD inhibition to what we believe is an attempt to regain cell homeostasis. One of these genes, *GADD45A,* is robustly increased by adipocyte exposure in SCD-insensitive cell lines and by SCD inhibition in SCD-sensitive cell lines. *GADD45A* is often induced by DNA damage and other stress signals [43] and more recently has been shown to sustain low levels of reactive oxygen species, thereby safeguarding leukemia cells for self-renewal [44]. Therefore, *GADD45A* may help mitigate the high LPO levels in PC3 cells, while also sustaining the activation of DNA damage and repair pathways, likely induced during adipocyte exposure by high LPO levels in SCD inhibition-insensitive cells. The induction of *GADD45A* and ER stress response, as evidenced by increased ATF4 protein expression, may enable these cells to adapt to oxidative stress in the harsh bone marrow microenvironment independently of SCD activity, rendering them less sensitive to SCD inhibition.

One notable characteristic of PC3 cells, which exhibit low sensitivity to SCD inhibition, is their high expression of lipid transporters *FABP4* and *CD36*, along with increased uptake of adipocyte-derived lipids. Unlike SCD-sensitive ARCaP(M) cells, PC3 cells do not show increased DNA damage or decreased mTOR signaling following SCD inhibition. Instead, they further increase the expression of *FABP4* and *CD36*, proteins shown to promote cancer progression by activating the PI3K/mTOR pathway [31-33]. This suggests that the enhanced lipid uptake and transport may be helping sustain mTOR signaling in PC3 cells, thereby preventing the anti-tumor effects observed in SCD-sensitive cell lines. It should be noted that our studies also demonstrated that, although direct inhibition of mTOR signaling with Everolimus significantly reduces PC3 spheroid growth, combining it with SCD deficiency does not produce an additive or synergistic effect. These results suggest that SCD-insensitive cell lines, with higher expression of *CD36* and *FAPB4,* depend on mTOR signaling for survival and maintenance of redox balance, but cannot be sensitized to SCD inhibition via mTOR blockade. This further indicates that intrinsic characteristics or dependencies of cancer cells may underlie their sensitivity or resistance to SCD inhibition, and these attributes cannot always be artificially induced by targeting the downstream pathways.

Prior studies by others have demonstrated that oncogenic activation of the PI3K/AKT/mTOR pathway confers resistance to ferroptotic cell death through SCD activity [45]. Our results align with these data and demonstrate that SCD reduces phosphorylation of AKT and downstream mTOR proteins, but only in PCa cell lines that do not harbor PTEN genomic aberrations (ARCaP(M), 22RV1) and have low lipid transporter expression and uptake. PTEN loss is one of the most common genomic aberrations in prostate cancer [46], and is strongly associated with poor clinical outcomes [47]. Our study showed for the first time that SCD-sensitive cell lines, which have decreased lipid uptake and wild-type PTEN, show reduced DNA strand breaks during adipocyte exposure, a phenotype that is reversed by SCD inhibition. Notably, PTEN has been found to suppress fatty acid accumulation in goat mammary epithelial cells, and liver-specific knockout of PTEN was shown to result in triglyceride accumulation [48]. The increased expression of lipid uptake proteins *CD36* and *FABP4*, along with PTEN loss, might serve as a protective mechanism against DNA damage, rendering these cells less sensitive to SCD inhibition. Indeed, the formation of lipid droplets has been demonstrated in cancer cells, shielding them from oxaliplatin-induced DNA damage [49]. Although SCD inhibition has been previously linked to DNA damage, particularly in the context of temozolomide resistance in glioblastoma [50, 51], no studies to date have suggested that the induction of DNA damage during SCD inhibition could be driven by increased lipid transporter expression, enhanced lipid uptake, and genetic aberrations in PTEN.

Bone metastatic PCa typically shows resistance to DNA-damaging agents such as platinum-based therapies or PARP inhibitors when used as monotherapies [52]. Our results demonstrate that SCD inhibition in SCD-sensitive cell lines interacting with adipocytes leads to upregulation of several genes within the HRR pathway. This suggests that targeting lipid metabolism in combination with DNA-damaging agents could be a promising therapeutic strategy for bone metastatic PCa. One potential candidate for combination therapy with SCD inhibitors is the PSMA-specific, beta particle-emitting peptidomimetic lutetium-177 (^177^Lu-PSMA-617). Given that introducing DNA strand breaks is the primary mechanism of ^177^Lu-PSMA-617 cytotoxicity and that improved overall survival with ^177^Lu-PSMA-617 has been reported in metastatic PCa patients who progressed on taxane therapy [53, 54], combining this with SCD inhibition could significantly enhance tumor response, which warrants further investigations. Another potential strategy is to combine SCD inhibition with an inhibitor of Ataxia telangiectasia mutated (ATM), particularly in tumors that are PTEN-deficient or harbor PTEN mutations. Loss of PTEN has been associated with increased oxidative stress, a greater reliance on ATM for redox homeostasis, and heightened sensitivity to ATM inhibition [55]. Given our finding that PTEN-null cells (PC3, C42B) show limited sensitivity to SCD inhibition alone, the combination with an ATM inhibitor may offer a more effective therapeutic approach.

We acknowledge several limitations in our *in vivo* study. First, we only performed SCD inhibition studies in mice fed HFD. We selected the HFD model of marrow adiposity based on its relevance to our *in vitro* findings, which demonstrated a cell type-dependent effect of marrow adipocytes on SCD expression and sensitivity to SCD inhibition. Additionally, our studies were conducted in immunocompromised mice, which prevented us from assessing the immune response to SCD inhibition. Given the role of SCD in regulatory T-cell differentiation [56], targeting the immune microenvironment in bone may potentially enhance the anti-tumor response during SCD inhibition, which will be a subject of future studies. Lastly, CAY10566, used in our study to pharmacologically inhibit SCD activity, has shown some undesired side effects in pre-clinical models [57]. However, newer inhibitors, such as SSI-4 [58], have demonstrated better safety profiles *in vivo* and are expected to move into the clinical setting. Nonetheless, the toxicity of current SCD pharmacological inhibitors remains a significant challenge.

Studies presented herein identify SCD as a modulator of cellular stress and a promising molecular target in metastatic PCa. We show that SCD inhibition in sensitive cell lines results in lipid-induced stress that leads to DNA damage, ER stress, and reduced survival. Importantly, we also reveal that lipid uptake and transport mechanisms, genomic aberrations in DDR genes and PTEN status may influence tumor response to SCD inhibition, highlighting the need for further investigation. Collectively, the results from our study underscore the role of SCD as a key metabolic regulator of adipocyte-tumor crosstalk in the bone metastatic niche and emphasize the importance of exploring the molecular mechanisms underlying tumor cell resilience and survival in the lipid-rich bone marrow niche.

## ACKNOWLEDGEMENTS

This research was supported in part by NIH grants R01CA251394 (IP), T32CA009531 (AW), F31CA284576 (AW) and P30CA22453 (MICR). Pilot funds were provided by the Karmanos Cancer Institute (KCI SRIG) (IP). The Microscopy, Imaging and Cytometry Resources Core was supported, in part by NIH Center grants P30 CA22453 to the Karmanos Cancer Institute and R50 CA251068-01 to Dr. Moin, Wayne State University.

## AUTHOR CONTRIBUTIONS

**Alexis Wilson:** conceptualization, methodology, investigation, validation, formal analysis, visualization, writing – Original draft preparation and Review & Editing; **Mackenzie K Herroon**: investigation, validation, writing – Review & Editing; **Shane Mecca**: investigation, formal analysis, writing – Review & Editing;: investigation, **Laimar Garmo:** investigation, writing – Review & Editing; **Jacob Lindquist:** investigation, writing – Review & Editing; **Shrila Rajendran:** investigation, formal analysis, writing – Review & Editing; **Steve M Patrick**: investigation, resources, writing – Review & Editing, **Izabela Podgorski:** conceptualization, methodology, formal analysis, resources, writing – Original draft preparation and Review & Editing; supervision, funding acquisition.

## REFERENCES

1. Ackerman D & Simon MC (2014/08/01) Hypoxia, lipids, and cancer: surviving the harsh tumor microenvironment. Trends in Cell Biology 24, doi: 10.1016/j.tcb.2014.06.001.

2. Scott JS, Nassar ZD, Swinnen JV & Butler LM (2022) Monounsaturated Fatty Acids: Key Regulators of Cell Viability and Intracellular Signaling in Cancer. Molecular Cancer Research 20, 1354–1364, doi: 10.1158/1541-7786.mcr-21-1069.

3. Tracz-Gaszewska Z & Dobrzyn P (2019) Stearoyl-CoA Desaturase 1 as a Therapeutic Target for the Treatment of Cancer. Cancers 11, 948, doi: 10.3390/cancers11070948.

4. Diedrich JD, Herroon MK, Rajagurubandara E & Podgorski I (2018) The Lipid Side of Bone Marrow Adipocytes: How Tumor Cells Adapt and Survive in Bone. Current Osteoporosis Reports 16, 443–457, doi: 10.1007/s11914-018-0453-9.

5. Sen U, Coleman C & Sen T (2023/06/01) Stearoyl coenzyme A desaturase-1: multitasker in cancer, metabolism, and ferroptosis. Trends in Cancer 9, doi: 10.1016/j.trecan.2023.03.003.

6. Igal RA (2011) Roles of StearoylCoA Desaturase-1 in the Regulation of Cancer Cell Growth, Survival and Tumorigenesis. Cancers 3, 2462–2477, doi: 10.3390/cancers3022462.

7. Herroon MK, Diedrich JD, Rajagurubandara E, Martin C, Maddipati KR, Kim S, Heath EI, Granneman J & Podgorski I (2019) Prostate Tumor Cell–Derived IL1β Induces an Inflammatory Phenotype in Bone Marrow Adipocytes and Reduces Sensitivity to Docetaxel via Lipolysis-Dependent Mechanisms. Molecular Cancer Research 17, 2508–2521, doi: 10.1158/1541-7786.mcr-19-0540.

8. Herroon MK, Rajagurubandara E, Hardaway AL, Powell K, Turchick A, Feldmann D & Podgorski I (2013/11) Bone marrow adipocytes promote tumor growth in bone via FABP4-dependent mechanisms. Oncotarget 4, doi: 10.18632/oncotarget.1482.

9. Ladanyi A, Mukherjee A, Kenny HA, Johnson A, Mitra AK, Sundaresan S, Nieman KM, Pascual G, Benitah SA, Montag A, Yamada SD, Abumrad NA & Lengyel E (2018) Adipocyte-induced CD36 expression drives ovarian cancer progression and metastasis. Oncogene 37, 2285–2301, doi: 10.1038/s41388-017-0093-z.

10. Shafat MS, Oellerich T, Mohr S, Robinson SD, Edwards DR, Marlein CR, Piddock RE, Fenech M, Zaitseva L, Abdul-Aziz A, Turner J, Watkins JA, Lawes M, Bowles KM & Rushworth SA (2017) Leukemic blasts program bone marrow adipocytes to generate a protumoral microenvironment. Blood 129, 1320–1332, doi: 10.1182/blood-2016-08-734798.

11. Herroon MK, Mecca S, Haimbaugh A, Garmo LC, Rajagurubandara E, Todi SV, Baker TR & Podgorski I (2021) Adipocyte-driven unfolded protein response is a shared transcriptomic signature of metastatic prostate carcinoma cells. Biochimica et Biophysica Acta (BBA) - Molecular Cell Research 1868, 119101, doi: 10.1016/j.bbamcr.2021.119101.

12. Herroon MK, Rajagurubandara E, Diedrich JD, Heath EI & Podgorski I (2018) Adipocyte-activated oxidative and ER stress pathways promote tumor survival in bone via upregulation of Heme Oxygenase 1 and Survivin. Scientific Reports 8, doi: 10.1038/s41598-017-17800-5.

13. Podgorski I, Linebaugh BE, Koblinski JE, Rudy DL, Herroon MK, Olive MB & Sloane BF (2009) Bone marrow-derived cathepsin K cleaves SPARC in bone metastasis. Am J Pathol 175, 1255–1269, doi: ajpath.2009.080906 [pii] 10.2353/ajpath.2009.080906.

14. Diedrich JD, Rajagurubandara E, Herroon MK, Mahapatra G, Hüttemann M & Podgorski I (2016) Bone marrow adipocytes promote the Warburg phenotype in metastatic prostate tumors via HIF-1α activation. Oncotarget 7, 64854–64877, doi: 10.18632/oncotarget.11712.

15. Heyza JR, Arora S, Zhang H, Conner KL, Lei W, Floyd AM, Deshmukh RR, Sarver J, Trabbic CJ, Erhardt P, Chan T-H, Dou QP & Patrick SM (2018/11) Targeting the DNA Repair Endonuclease ERCC1-XPF with Green Tea Polyphenol Epigallocatechin-3-Gallate (EGCG) and Its Prodrug to Enhance Cisplatin Efficacy in Human Cancer Cells. Nutrients 10, doi: 10.3390/nu10111644.

16. Herroon MK, Diedrich JD & Podgorski I (2016) New 3D-Culture Approaches to Study Interactions of Bone Marrow Adipocytes with Metastatic Prostate Cancer Cells. Frontiers in Endocrinology 7, doi: 10.3389/fendo.2016.00084.

17. Dobin A, Davis CA, Schlesinger F, Drenkow J, Zaleski C, Jha S, Batut P, Chaisson M & Gingeras TR (2013) STAR: ultrafast universal RNA-seq aligner. Bioinformatics 29, 15–21, doi: 10.1093/bioinformatics/bts635.

18. Chen Y, Chen L, Lun Aaron TL, Baldoni Pedro L & Smyth Gordon K (2025) edgeR v4: powerful differential analysis of sequencing data with expanded functionality and improved support for small counts and larger datasets. Nucleic Acids Research 53, doi: 10.1093/nar/gkaf018.

19. Liberzon A, Birger C, Thorvaldsdóttir H, Ghandi M, Mesirov Jill P & Tamayo P (2015) The Molecular Signatures Database Hallmark Gene Set Collection. Cell Systems 1, 417–425, doi: 10.1016/j.cels.2015.12.004.

20. Falank C, Fairfield H & Reagan MR (2016) Signaling Interplay between Bone Marrow Adipose Tissue and Multiple Myeloma cells. Front Endocrinol (Lausanne) 7, 67, doi: 10.3389/fendo.2016.00067.

21. Reagan MR, Fairfield H & Rosen CJ (2021) Bone Marrow Adipocytes: A Link between Obesity and Bone Cancer. Cancers (Basel) 13, doi: 10.3390/cancers13030364.

22. Collins JM, Neville MJ, Hoppa MB & Frayn KN (02/26/2010) De novo lipogenesis and stearoyl-CoA desaturase are coordinately regulated in the human adipocyte and protect against palmitate-induced cell injury - PubMed. The Journal of biological chemistry 285, doi: 10.1074/jbc.M109.053280.

23. Collins JM, Neville MJ, Pinnick KE, Hodson L, Ruyter B, van Dijk TH, Reijngoud D-J, Fielding MD & Frayn KN (2011 Sep) De novo lipogenesis in the differentiating human adipocyte can provide all fatty acids necessary for maturation - PubMed. Journal of lipid research 52, doi: 10.1194/jlr.M012195.

24. Bian X, Liu R, Meng Y, Xing D, Xu D & Lu Z (2021/01/01) Cancer Focus: Lipid metabolism and cancer. The Journal of Experimental Medicine 218, doi: 10.1084/jem.20201606.

25. Martin-Perez M, Urdiroz-Urricelqui U, Bigas C & Benitah SA (2022) The role of lipids in cancer progression and metastasis. Cell Metab 34, 1675–1699, doi: 10.1016/j.cmet.2022.09.023.

26. Flor AC, Wolfgeher D, Wu D, Kron SJ, Flor AC, Wolfgeher D, Wu D & Kron SJ (2017-10-30) A signature of enhanced lipid metabolism, lipid peroxidation and aldehyde stress in therapy-induced senescence. Cell Death Discovery 2017 3:1 3, doi: 10.1038/cddiscovery.2017.75.

27. Kawaguchi Y, Hagiwara D, Miyata T, Hodai Y, Kurimoto J, Takagi H, Suga H, Kobayashi T, Sugiyama M, Onoue T, Ito Y, Iwama S, Banno R, Grinevich V, Arima H, Kawaguchi Y, Hagiwara D, Miyata T, Hodai Y, Kurimoto J, Takagi H, Suga H, Kobayashi T, Sugiyama M, Onoue T, Ito Y, Iwama S, Banno R, Grinevich V & Arima H (2020-11-12) Endoplasmic reticulum chaperone BiP/GRP78 knockdown leads to autophagy and cell death of arginine vasopressin neurons in mice. Scientific Reports 2020 10:1 10, doi: 10.1038/s41598-020-76839-z.

28. Williams B, Verchot J & Dickman MB (2014/06/04) Frontiers | When supply does not meet demand-ER stress and plant programmed cell death. Frontiers in Plant Science 5, doi: 10.3389/fpls.2014.00211.

29. Zou Z, Tao T, Li H & Zhu X (2020) mTOR signaling pathway and mTOR inhibitors in cancer: progress and challenges. Cell & Bioscience 10, doi: 10.1186/s13578-020-00396-1.

30. Herroon MK, Rajagurubandara E, Hardaway AL, Powell K, Turchick A, Feldmann D & Podgorski I (2013) Bone marrow adipocytes promote tumor growth in bone via FABP4-dependent mechanisms. Oncotarget 4, 2108–2123, doi: 10.18632/oncotarget.1482.

31. Gao Y, Wang Y, Wang X, Ma J, Wei M, Li N & Zhao Z (2023 Jun) FABP4 Regulates Cell Proliferation, Stemness, Apoptosis, and Glycolysis in Colorectal Cancer via Modulating ROS/ERK/mTOR Pathway - PubMed. Discovery medicine 35, doi: 10.24976/Discov.Med.202335176.37.

32. Liu H, Guo W, Wang T, Cao P, Zou T, Peng Y, Yan T, Liao C, Li Q, Duan Y, Han J, Zhang B, Chen Y, Zhao D & Yang X (2024 Feb 6) CD36 inhibition reduces non-small-cell lung cancer development through AKT-mTOR pathway. Cell Biology and Toxicology 40, doi: 10.1007/s10565-024-09848-7.

33. Luo X, Zheng E, Wei L, Zeng H, Qin H, Zhang X, Liao M, Chen L, Zhao L, Ruan XZ, Yang P, Chen Y, Luo X, Zheng E, Wei L, Zeng H, Qin H, Zhang X, Liao M, Chen L, Zhao L, Ruan XZ, Yang P & Chen Y (2021-03-26) The fatty acid receptor CD36 promotes HCC progression through activating Src/PI3K/AKT axis-dependent aerobic glycolysis. Cell Death & Disease 2021 12:4 12, doi: 10.1038/s41419-021-03596-w.

34. Dan HC, Ebbs A, Pasparakis M, Van Dyke T, Basseres DS & Baldwin AS (2014) Akt-dependent Activation of mTORC1 Complex Involves Phosphorylation of mTOR (Mammalian Target of Rapamycin) by IκB Kinase α (IKKα). Journal of Biological Chemistry 289, 25227–25240, doi: 10.1074/jbc.m114.554881.

35. Falletti O, Cadet J, Favier A & Douki T (2007) Trapping of 4-hydroxynonenal by glutathione efficiently prevents formation of DNA adducts in human cells. Free Radic Biol Med 42, 1258–1269, doi: 10.1016/j.freeradbiomed.2007.01.024.

36. Gentile F, Arcaro A, Pizzimenti S, Daga M, Cetrangolo GP, Dianzani C, Lepore A, Graf M, Ames PRJ & Barrera G (2017) DNA damage by lipid peroxidation products: implications in cancer, inflammation and autoimmunity. AIMS Genetics 4, doi: 10.3934/genet.2017.2.103.

37. Shoeb M, Ansari NH, Srivastava SK & Ramana KV (2014) 4-hydroxynonenal in the pathogenesis and progression of human diseases. Current medicinal chemistry 21, doi: 10.2174/09298673113209990181.

38. Barrera G (2012) Oxidative Stress and Lipid Peroxidation Products in Cancer Progression and Therapy. ISRN Oncology 2012, doi: 10.5402/2012/137289.

39. Pope LE & Dixon SJ (2023) Regulation of ferroptosis by lipid metabolism. Trends in Cell Biology 33, 1077–1087, doi: 10.1016/j.tcb.2023.05.003.

40. Tesfay L, Paul BT, Konstorum A, Deng Z, Cox AO, Lee J, Furdui CM, Hegde P, Torti FM & Torti SV (2019/10/15) Stearoyl-CoA Desaturase 1 Protects Ovarian Cancer Cells from Ferroptotic Cell Death. Cancer Research 79, doi: 10.1158/0008-5472.CAN-19-0369.

41. Krusenstiern ANv, Robson RN, Qian N, Qiu B, Hu F, Reznik E, Smith N, Zandkarimi F, Estes VM, Dupont M, Hirschhorn T, Shchepinov MS, Min W, Woerpel KA & Stockwell BR (2023/06) Identification of essential sites of lipid peroxidation in ferroptosis. Nature chemical biology 19, doi: 10.1038/s41589-022-01249-3.

42. Das UN (2019) Saturated Fatty Acids, MUFAs and PUFAs Regulate Ferroptosis. Cell Chem Biol 26, 309–311, doi: 10.1016/j.chembiol.2019.03.001.

43. Liebermann DA & Hoffman B (2008) Gadd45 in stress signaling. Journal of Molecular Signaling 3, 15, doi: 10.1186/1750-2187-3-15.

44. Hassan N, Yi H, Malik B, Gaspard-Boulinc L, Samaraweera SE, Casolari DA, Seneviratne J, Balachandran A, Chew T, Duly A, Carter DR, Cheung BB, Norris M, Haber M, Kavallaris M, Marshall GM, Zhang XD, Liu T, Wang J, Liebermann DA, D’Andrea RJ & Wang JY (2024) Loss of the stress sensor GADD45A promotes stem cell activity and ferroptosis resistance in LGR4/HOXA9-dependent AML. Blood 144, 84–98, doi: 10.1182/blood.2024024072.

45. Yi J, Zhu J, Wu J, Thompson CB & Jiang X (2020) Oncogenic activation of PI3K-AKT-mTOR signaling suppresses ferroptosis via SREBP-mediated lipogenesis. Proc Natl Acad Sci U S A 117, 31189–31197, doi: 10.1073/pnas.2017152117.

46. Hamid AA, Gray KP, Shaw G, MacConaill LE, Evan C, Bernard B, Loda M, Corcoran NM, Van Allen EM, Choudhury AD & Sweeney CJ (2019) Compound Genomic Alterations of TP53, PTEN, and RB1 Tumor Suppressors in Localized and Metastatic Prostate Cancer. Eur Urol 76, 89–97, doi: 10.1016/j.eururo.2018.11.045.

47. Jamaspishvili T, Berman DM, Ross AE, Scher HI, Marzo AMD, Squire JA & Lotan TL (2018/04) Clinical implications of PTEN loss in prostate cancer. Nature reviews Urology 15, doi: 10.1038/nrurol.2018.9.

48. Yao DW, Ma J, Yang CL, Chen LL, He QY, Coleman DN, Wang TZ, Jiang XL, Luo J, Ma Y & Loor JJ (2021/06/01) Phosphatase and tensin homolog (PTEN) suppresses triacylglycerol accumulation and monounsaturated fatty acid synthesis in goat mammary epithelial cells. Journal of Dairy Science 104, doi: 10.3168/jds.2020-18784.

49. Yu J, Hu D, Cheng Y, Guo J, Wang Y, Tan Z, Peng J & Zhou H (2021/06/05) Lipidomics and transcriptomics analyses of altered lipid species and pathways in oxaliplatin-treated colorectal cancer cells. Journal of Pharmaceutical and Biomedical Analysis 200, doi: 10.1016/j.jpba.2021.114077.

50. Dai S, Yan Y, Xu Z, Zeng S, Qian L, Huo L, Li X, Sun L & Gong Z (2017) SCD1 Confers Temozolomide Resistance to Human Glioma Cells via the Akt/GSK3β/β-Catenin Signaling Axis. Front Pharmacol 8, 960, doi: 10.3389/fphar.2017.00960.

51. Eyme KM, Sammarco A, Jha R, Mnatsakanyan H, Pechdimaljian C, Carvalho L, Neustadt R, Moses C, Alnasser A, Tardiff DF, Su B, Williams KJ, Bensinger SJ, Chung CY & Badr CE (2023) Targeting de novo lipid synthesis induces lipotoxicity and impairs DNA damage repair in glioblastoma mouse models. Science Translational Medicine 15, eabq6288, doi: doi:10.1126/scitranslmed.abq6288.

52. Hager S, Ackermann CJ, Joerger M, Gillessen S & Omlin A (2016) Anti-tumour activity of platinum compounds in advanced prostate cancer&#x2014;a systematic literature review. Annals of Oncology 27, 975–984, doi: 10.1093/annonc/mdw156.

53. Bauckneht M, Ciccarese C, Laudicella R, Mosillo C, D’Amico F, Anghelone A, Strusi A, Beccia V, Bracarda S, Fornarini G, Tortora G & Iacovelli R (2024) Theranostics revolution in prostate cancer: Basics, clinical applications, open issues and future perspectives. Cancer Treat Rev 124, 102698, doi: 10.1016/j.ctrv.2024.102698.

54. Stracker TH, Osagie OI, Escorcia FE & Citrin DE (2023) Exploiting the DNA Damage Response for Prostate Cancer Therapy. Cancers (Basel) 16, doi: 10.3390/cancers16010083.

55. McCabe N, Hanna C, Walker SM, Gonda D, Li J, Wikstrom K, Savage KI, Butterworth KT, Chen C, Harkin DP, Prise KM & Kennedy RD (2015) Mechanistic Rationale to Target PTEN-Deficient Tumor Cells with Inhibitors of the DNA Damage Response Kinase ATM. Cancer Res 75, 2159–2165, doi: 10.1158/0008-5472.Can-14-3502.

56. Grajchen E, Loix M, Baeten P, Côrte-Real BF, Hamad I, Vanherle S, Haidar M, Dehairs J, Broos JY, Ntambi JM, Zimmermann R, Breinbauer R, Stinissen P, Hellings N, Verberk SGS, Kooij G, Giera M, Swinnen JV, Broux B, Kleinewietfeld M, Hendriks JJA & Bogie JFJ (2023) Fatty acid desaturation by stearoyl-CoA desaturase-1 controls regulatory T cell differentiation and autoimmunity. Cellular & Molecular Immunology 20, 666–679, doi: 10.1038/s41423-023-01011-2.

57. Zhang Z, A. Dn & D. Wm (2014) Opportunities and Challenges in Developing Stearoyl-Coenzyme A Desaturase-1 Inhibitors as Novel Therapeutics for Human Disease. Journal of Medicinal Chemistry 57, doi: 10.1021/jm401516c.

58. Sun S, Zhang Z, Pokrovskaia N, Chowdhury S, Jia Q, Chang E, Khakh K, Kwan R, McLaren DG, Radomski CC, Ratkay LG, Fu J, Dales NA & Winther MD (2015) Discovery of triazolone derivatives as novel, potent stearoyl-CoA desaturase-1 (SCD1) inhibitors. Bioorg Med Chem 23, 455–465, doi: 10.1016/j.bmc.2014.12.014.

