## Supplementary Material for "Targeting SCD in SCD-amplified prostate cancer inhibits growth in bone by modulating cellular stress, mTOR, and DNA damage pathways"

<u>**Supplemental Figures**</u>

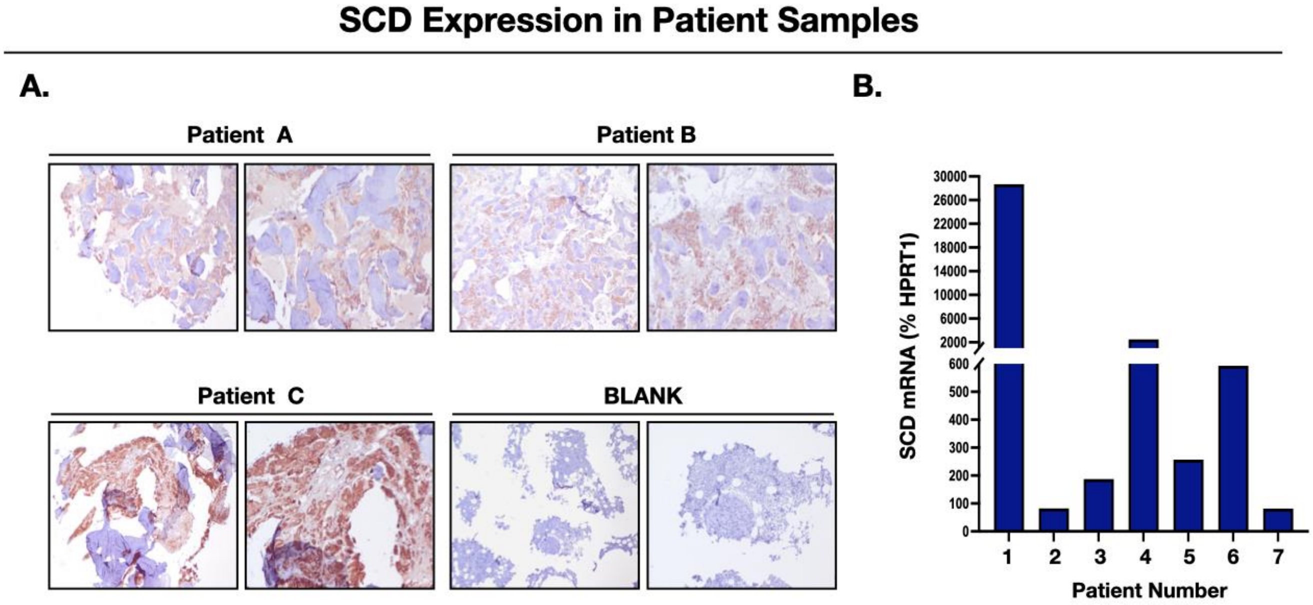


**Supplementary Figure 1. SCD is expressed in bone metastatic samples from PCa patients. A:** Immunohistochemical (NovaRed) analysis of SCD expression in bone metastatic lesions from PCa patients. 20× images; "Blank" panels indicate no primary antibody control. **B:** *SCD* mRNA levels in metastatic bone lesions from PCa patients; Relative levels were calculated using $2^{-\Delta Ct}$. Results for each patient are shown as percent housekeeping gene (*HPRT1*).

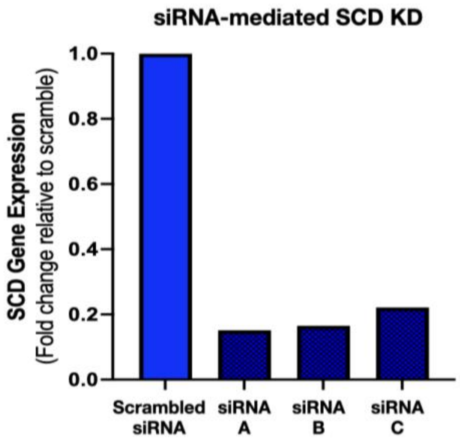


**Supplementary Figure 2. SCD knockdown by individual siRNA duplexes. A:** *SCD* mRNA levels in ARCaP(M) cells upon treatment with nonoverlapping siRNAs targeting SCD as determined by TaqMan RT-PCR analysis.

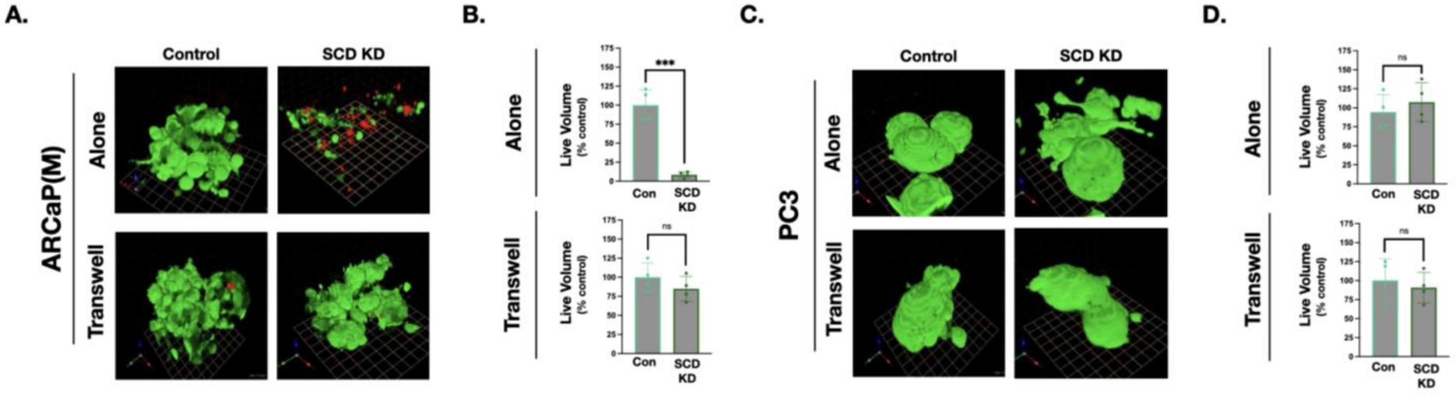


**Supplementary Figure 3. Knockdown of SCD reduces the size of ARCaP(M) spheroids.** ARCaP(M) **(A, B)** and PC3 **(C, D)** cells were grown in 3-dimensional cultures on reconstituted basement membrane (rBM) with a 2% Cultrex overlay. Live/Dead assay on 3D cultures grown alone **(top panels)** or with adipocytes **(bottom panels)** and treated with scrambled siRNA (Control) or siRNA targeting SCD. Live cells (green fluorescence; calcein AM); dead cells (red fluorescence; ethidium homodimer). **B, D**: Quantification of live (green) spheroid volume compared to control for ARCaP(M) **(B)** and PC3 cells **(D)** grown alone **(top panel)** or with adipocytes **(bottom panel)**; *** p < 0.001, ns = not significant.

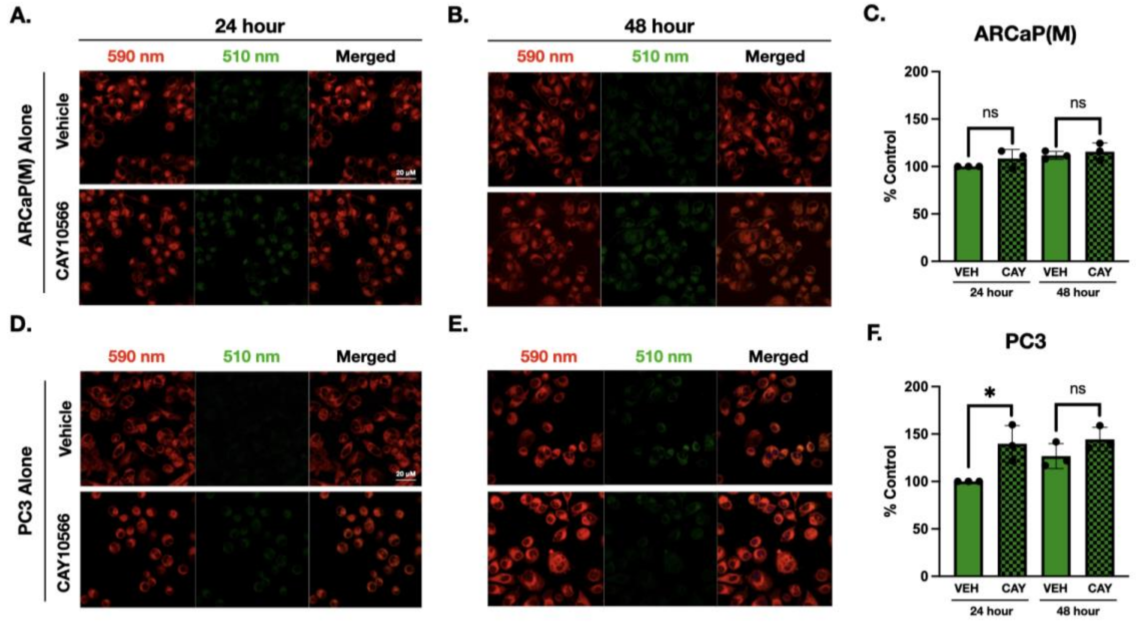


**Supplementary Figure 4. SCD pharmacological inhibition does not increase lipid peroxidation in PCa cells grown in the absence of adipocytes.** ARCaP(M) **(A-C)** and PC3 **(D-F)** cells were grown in alone conditions and treated with vehicle control (0.1% DMSO) or 1µM CAY10566 for 24 **(A, D)** and 48 **(B, E)** hours. BODIPY C11 staining was performed to examine LPO levels. An increase in LPO is indicated by a shift of fluorescence emission peak from ~590 nm (red) towards ~510 nm (green); 40x images. Quantification of ~510 nm shift for ARCaP(M) cells **(C)** and PC3 cells **(F).** * p < 0.05, ns = not significant.

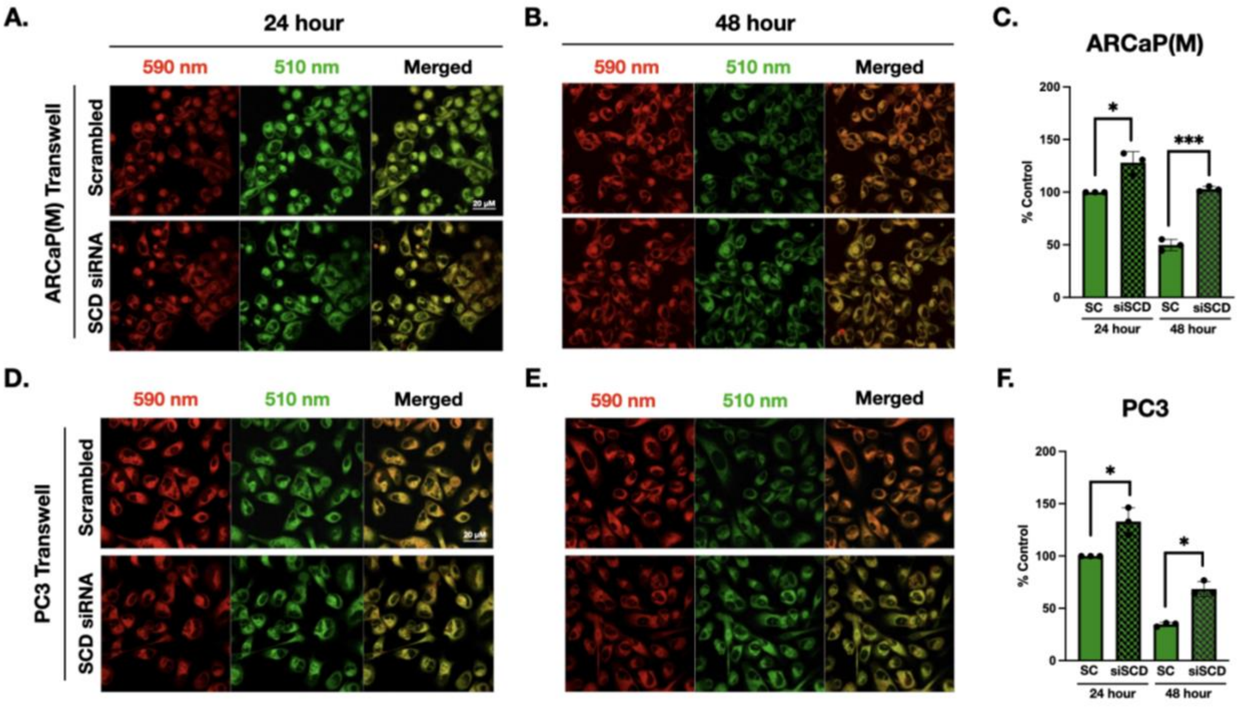


**Supplementary Figure 5. siRNA-mediated SCD knockdown increases lipid peroxidation levels in PCa cells in Transwell co-culture with adipocytes.** ARCaP(M) **(A-C)** and PC3 **(D-F)** cells were treated with scrambled siRNA or siRNA targeting SCD and grown in TW co-culture with adipocytes for 24 **(A, D)** and 48 **(B, E)** hours. BODIPY C11 staining was performed to examine LPO levels. An increase in LPO is indicated by a shift of fluorescence emission peak from ~590 nm (red) towards ~510 nm (green); 40x images. Quantification of ~510 nm shift for ARCaP(M) **(C)** and PC3 cells **(F),** indicating an increase in LPO upon treatment with siRNA targeting SCD as compared to scramble control; *$p < 0.05$, *** $p < 0.001$.

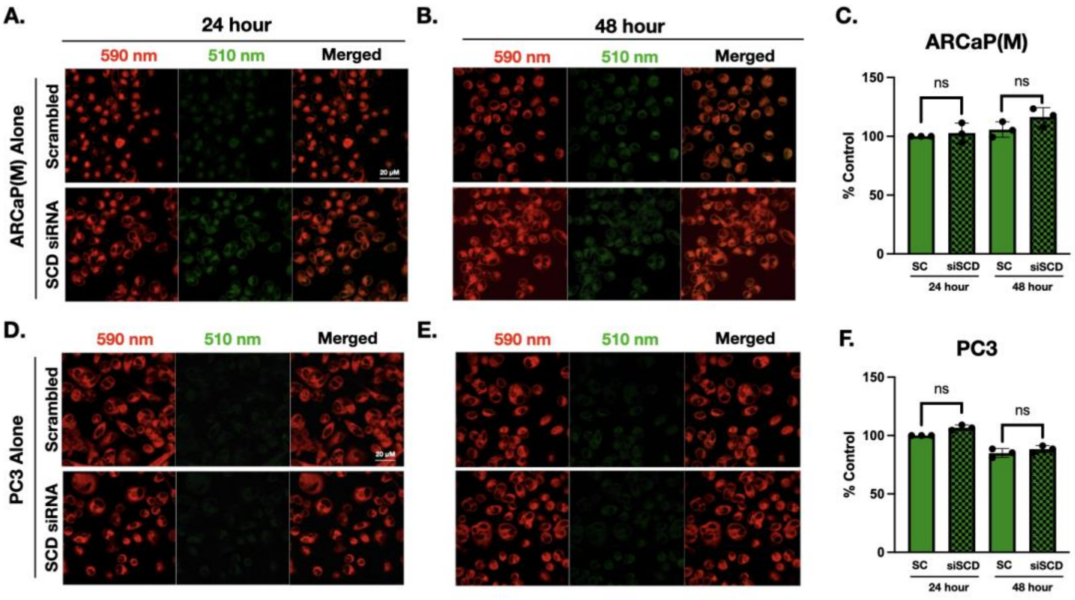


**Supplementary Figure 6. siRNA-mediated knockdown of SCD does not increase lipid peroxidation in PCa cells grown in the absence of adipocytes.** ARCaP(M) **(A-C)** and PC3 **(D-F)** cells were treated with scrambled siRNA or siRNA targeting SCD and grown in alone conditions for 24 **(A, D)** and 48 **(B, E)** hours. BODIPY C11 staining was performed to examine LPO levels. An increase in LPO is indicated by a shift of fluorescence emission peak from ~590 nm (red) towards ~510 nm (green); 40x images. Quantification of ~510 nm shift for ARCaP(M) cells **(C)** and PC3 cells **(F),** indicating no change in LPO upon treatment with siRNA targeting SCD as compared to scramble control; ns = not significant.

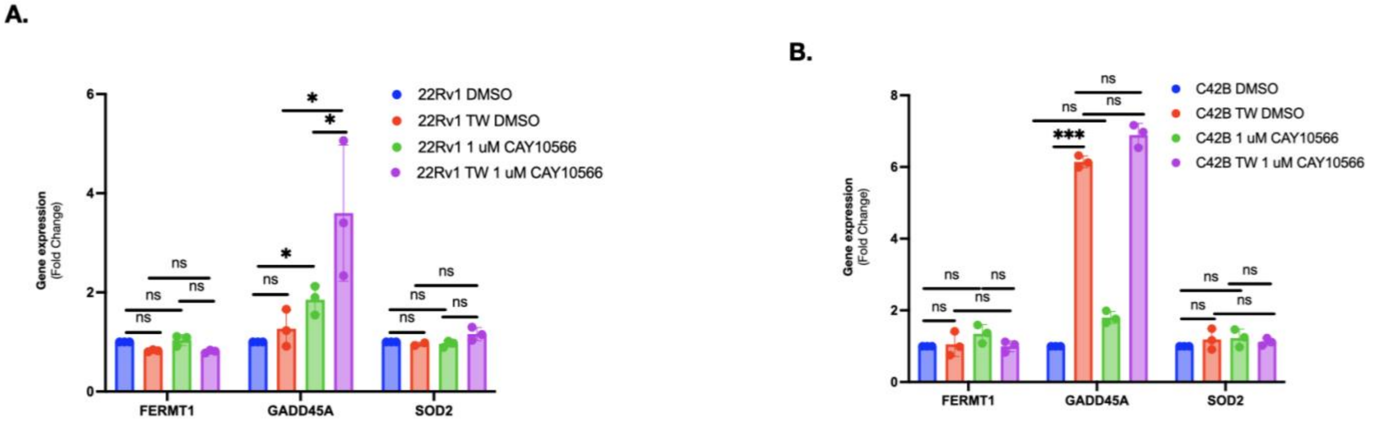


**Supplementary Figure 7. SCD inhibition induces expression of GADD45A in 22Rv1 cells only.** TaqMan RT-PCR analysis for mRNA expression of *FERMT1, GADD45A* and *SOD2* in 22Rv1 **(A)** and C42B **(B)** cells grown alone or in TW co-culture with adipocytes and treated with vehicle control (0.1% DMSO) or 1μM CAY10566. Data represent at least 3 experiments; *p < 0.05, *** p < 0.001, and ns = not significant.

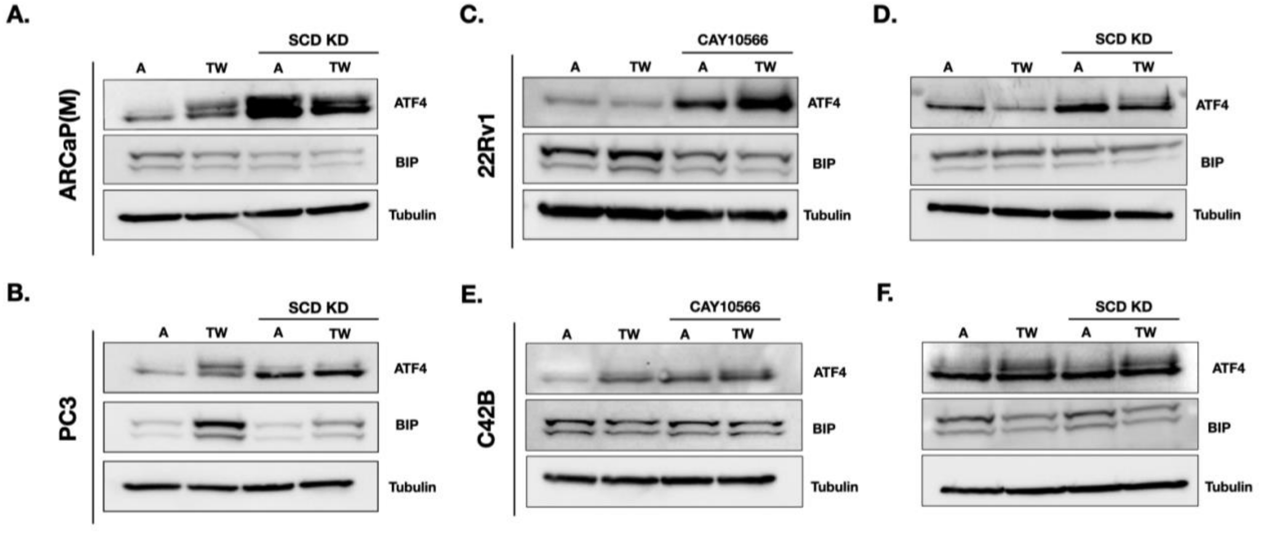


**Supplementary Figure 8. SCD knockdown/inhibition increases ER stress levels in PCa cells.** Immunoblot analysis of ER stress markers ATF4 (top) and BIP (bottom) in ARCaP(M) **(A)**, PC3 **(B)**, 22Rv1 **(C, D)** and C42B **(E, F)** cells grown alone or in TW co-culture with adipocytes and treated with vehicle control (0.1 % DMSO or scrambled siRNA), 1μM CAY10566 **(C, E)** or siRNA targeting SCD **(A, B, D, F).**

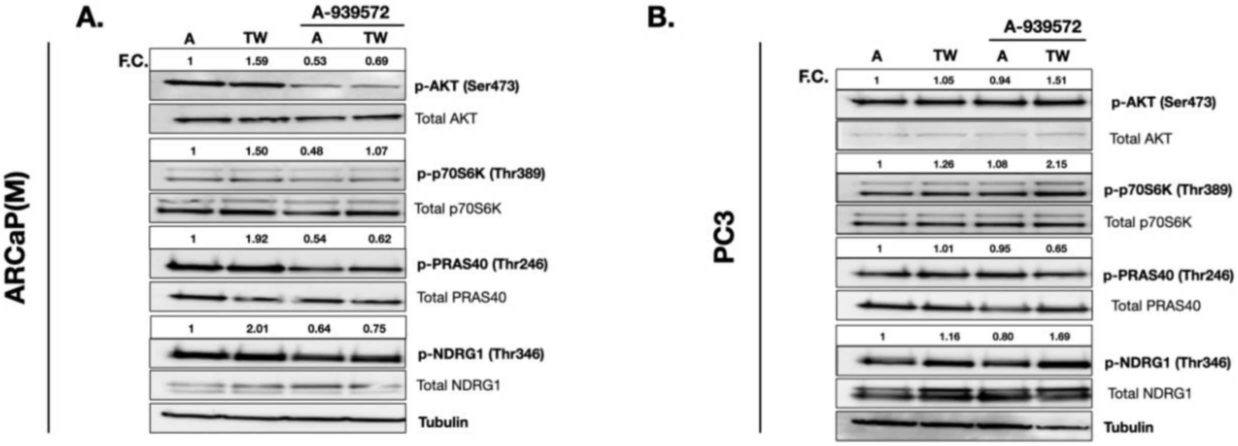


**Supplementary Figure 9. Inhibition of SCD decreases mTOR signaling in ARCaP(M) but not PC3 cells.** ARCaP(M) **(A)** and PC3 **(B)** cells were cells grown alone or in TW co-culture with adipocytes and treated with vehicle control (0.1% DMSO) or 250μM A-939572 for 48 hours and subjected to immunoblot analysis of total and phosphorylated downstream mTOR proteins: AKT, P70S6K, PRAS40 and NDRG1. Data represent at least 3 experiments.

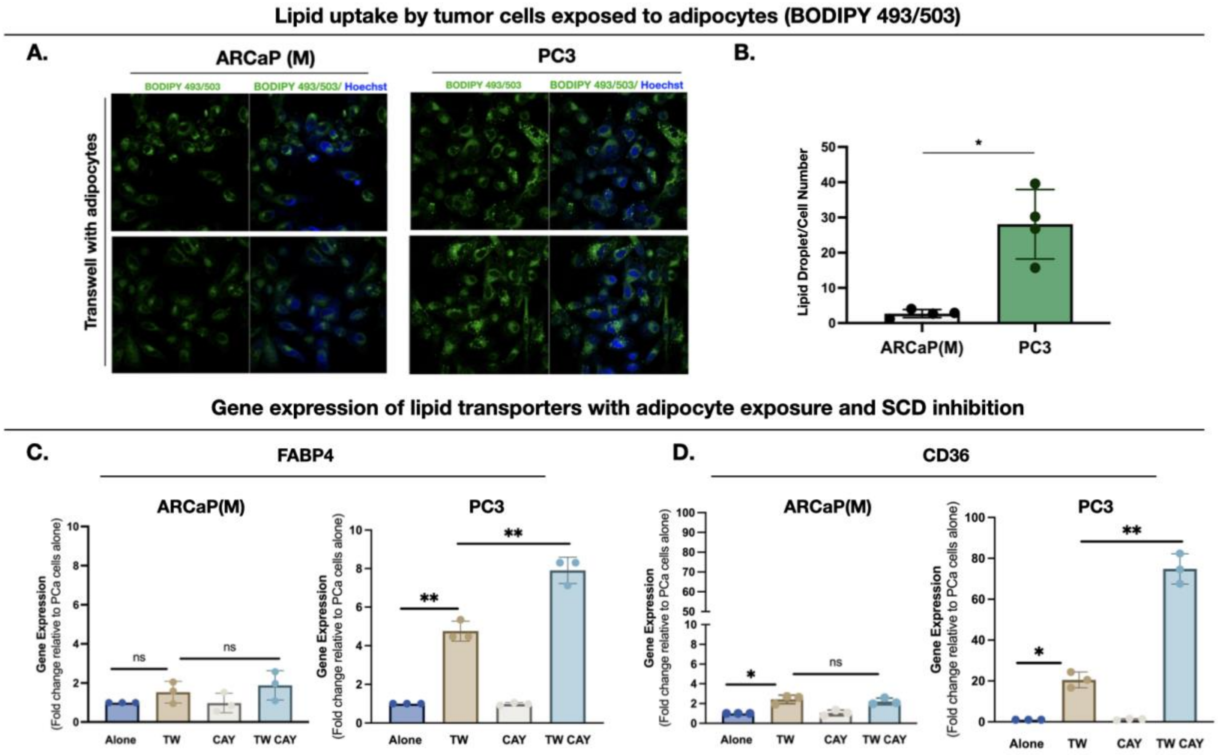


**Supplementary Figure 10. PC3 cells exposed to adipocytes have increased lipid uptake and expression of lipid transporters, which are further increased by SCD inhibition. A:** Immunofluorescence imaging of lipid droplets (BODIPY 493/503; green fluorescence) in ARCaP(M) and PC3 cells grown in TW co-culture with adipocytes for 48 hours; 40× images; Hoechst dye (blue) was used to stain the nuclei. **B:** Quantification of lipid droplet number per cell was determined using ImageJ's analyze particles function. **C-D:** TaqMan RT-PCR analysis of *FABP4* **(C)** and *CD36* **(D)** in ARCaP(M) and PC3 cells grown alone or in TW co-culture with adipocytes in the absence or presence of 1μM CAY10566 (CAY); *p < 0.05, ** p < 0.01 and ns = not significant.

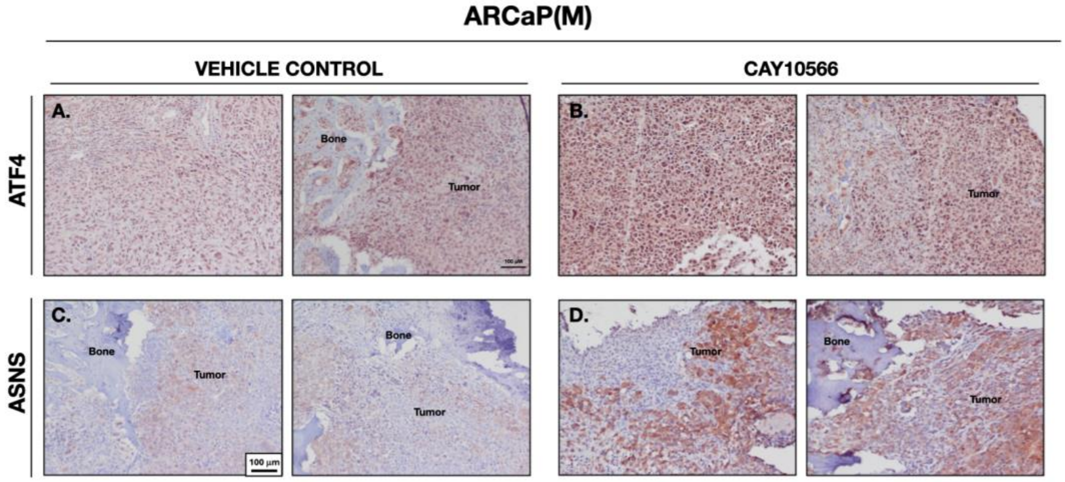


**Supplementary Figure 11. SCD inhibition *in vivo* increases ER stress and expression in ARCaP(M) bone tumors.** HFD-fed mice were intratibially implanted with ARCaP(M) cells and treated with vehicle control (5% DMSO in corn oil) or CAY10566 (5mg/kg) by oral gavage after confirmation of tumor formation. **A-D:** Immunohistochemical (NovaRed) staining of ATF4 **(A, B)** or ASNS **(C, D)** protein in bone tumors from vehicle-treated (**A, C**) and CAY10556-treated (**B, D**) mice; 10× images. Tumor, bone, and marrow are noted in images.
